# *Faecalibacterium prausnitzii*, depleted in the Parkinson’s disease microbiome, improves motor deficits in α-synuclein overexpressing mice

**DOI:** 10.1101/2025.09.18.677192

**Authors:** Anastasiya Moiseyenko, Giacomo Antonello, Aubrey M. Schonhoff, Joseph C. Boktor, Kaelyn Long, Blake Dirks, Anastasiya D. Oguienko, Alexander Viloria Winnett, Patrick Simpson, Dorsa Daeizadeh, Rustem F. Ismagilov, Rosa Krajmalnik-Brown, Nicola Segata, Levi D. Waldron, Sarkis K. Mazmanian

## Abstract

Gut microbiome composition is altered in Parkinson’s disease (PD), a neurodegenerative disorder characterized by motor dysfunction and frequently accompanied by gastrointestinal (GI) symptoms. Notably, microbial taxa with anti-inflammatory properties are consistently depleted in PD patients compared to controls. To explore whether specific gut bacteria may be disease-protective, we assembled a microbial consortium of 8 human-associated taxa that are reduced in individuals with PD across multiple cohorts and geographies. Treatment of α-synuclein overexpressing (Thy1-ASO) mice, an animal model of PD, with this consortium improved motor and GI deficits. A single bacterial species from this consortium, *Faecalibacterium prausnitzii*, was sufficient to correct gut microbiome deviations in Thy1-ASO mice, induce anti-inflammatory immune responses, and promote protective colonic gene expression profiles. Accordingly, oral treatment with *F. prausnitzii* robustly ameliorated motor and GI symptoms and reduced α-synuclein aggregates in the brain. These findings support the emerging hypothesis of functional contributions by the microbiome to PD and embolden development of potential probiotic therapies.

## Introduction

Parkinson’s disease (PD) is the second most common neurodegenerative disorder, affecting over 10 million people globally and 1% of individuals over the age of 60^1,2^. The diagnosis of PD is rapidly rising due to increased lifespan and potentially to certain environmental exposures^3^. Synucleinopathies such as PD, Lewy body dementia, and multiple system atrophy, among other disorders, are characterized by aggregation of the neuronal protein α-synuclein (αSyn)^4^. αSyn pathology can lead to inflammation, neuronal dysfunction, and likely to the ultimate death of dopaminergic neurons in the nigrostriatal pathway of the substantia nigra pars compacta (SNpc), resulting in motor symptoms such as tremors, rigidity, gait impairment, and bradykinesia^1,2,5^. Current treatments provide symptomatic relief, but can have considerable side effects, present dosing challenges, and often lose efficacy over time. There are no disease-modifying drugs for PD^2,6^.

Although PD is predominantly classified as a brain disorder, a role for non-motor symptoms and peripheral organs dates back over 200 years^7^. Indeed, 60-80% of PD patients experience gastrointestinal (GI) symptoms – primarily constipation, but also deficits in gastric emptying, abdominal pain, and increased intestinal permeability – that often appear many years before a PD diagnosis^8,9^. Other non-motor symptoms, including hyposmia, depression, and REM sleep behavior disorder, also manifest in the prodromal phase^10–12^. Braak’s hypothesis, and more recently the “brain-first vs. body-first” hypothesis^13,14^, postulates that αSyn aggregation may start in peripheral tissues such as the GI tract, spread via the vagus nerve, and eventually reach the brainstem, substantia nigra, and neocortex^15^. Indeed, injection of αSyn aggregates (as pre-formed fibrils [PFFs] or other pathogenic αSyn species) into the intestines of mice and rats results first in GI symptoms, and eventually leads to neurodegeneration in the brain and motor deficits^16,17^. Severing the vagus nerve in animals halts propagation of gut-induced αSyn pathology into the brain^16^ and epidemiologic data suggest that vagotomies are protective against development of PD in humans^18,19^. Whether PD can originate outside the brain, and potentially in the GI tract, is a timely yet still unresolved question.

Gut microbiome analyses of stool samples have revealed that individuals with PD harbor a distinct microbial profile compared to non-PD controls. PD patients show increased abundance of potential pathobionts and decreased levels of beneficial bacteria including *Blautia*, *Roseburia*, and *Faecalibacterium* species^20–24^. A recurring feature observed across numerous cohorts is loss of short-chain fatty acid (SCFA)-producing bacteria, many of which promote anti-inflammatory pathways^25–27^. While treatments containing Lactobacilli and Bifidobacteria have been shown to result in mild clinical improvements^28–31^, these taxa are among the most enriched in the PD patient microbiome^20^, raising questions about their therapeutic potential in the context of PD. Exploring the efficacy of microbes that are depleted in PD is a more intuitive therapeutic approach, yet remains largely unexplored.

Multiple metagenomic analyses have found *Faecalibacterium prausnitzii* to be consistently depleted from the gut microbiome of PD patients^20,21,24,32–36^. *F. prausnitzii*, a fastidiously anaerobic Gram-positive species, is highly abundant in the ileum and colon of the healthy human gut and is a major producer of the SCFA butyrate^37^. Its potent anti-inflammatory effects have been documented in models of inflammatory bowel disease (IBD)^37–40^. Notably, individuals with IBD show depletion of *F. prausnitzii* and are at increased risk for developing PD later in life^41,42^.

Using the transgenic Thy1 α-synuclein overexpressing (“Line 61”) mouse model of PD^43^ (hereafter referred to as Thy1-ASO), we report that treatment with a defined consortium of eight microbes commonly depleted in PD patients led to modest improvements in PD-like symptoms. One member of the consortium, *F. prausnitzii*, alone was able to robustly ameliorate motor and GI dysfunction and reduce αSyn accumulation in the brain. Additionally, *F. prausnitzii* treatment mildly remodeled the gut microbiome of Thy1-ASO mice to more closely resemble that of wild-type (WT) animals. Regulatory T cell (T_REGS_) numbers and production of the anti-inflammatory cytokine interleukin-10 (IL-10) were increased in *F. prausnitzii*-treated intestinal tissues. Furthermore, *F. prausnitzii* supplementation increased colonic expression of gene pathways associated with tissue repair and regeneration. Our findings demonstrate that treatment of Thy1-ASO mice with a commensal bacterium that is depleted in PD patients improves molecular and behavioral outcomes, providing conceptual validation for the development of microbial strategies to treat PD.

## Results

### A human gut microbial consortium ameliorates symptoms in Thy1-ASO mice

To investigate the potential therapeutic benefits of gut microbes, we used transgenic Thy1-ASO mice that overexpress the wildtype human *SNCA* gene under a modified Thy1 promoter, driving αSyn protein expression in many neuronal subtypes throughout the body and brain^43^. Thy1-ASO mice phenocopy clinical features of human synucleinopathies, including PD, such as αSyn pathology in the gut and brain, neuroinflammation, mitochondrial deficits, and progressive age-dependent deficits in motor and GI function^43,44^.

We assembled a defined microbial consortium composed of eight culturable type strains that represent the closest relatives of bacterial taxa depleted in stool samples across multiple human PD cohorts, which we termed “benCom-PD” for *<u>ben</u>*eficial *<u>com</u>*mensals for *<u>PD</u>*(**Fig. 1A**). BenCom-PD was intragastrically administered to specific pathogen-free (SPF; standard laboratory microbiome) Thy1-ASO mice twice weekly for approximately 15 weeks (**Fig. 1B**). At 20 weeks of age, motor and GI function was evaluated using a battery of standard behavioral tests. Mice receiving benCom-PD exhibited improvements in motor performance compared to vehicle-treated controls, with reduced latency in the adhesive removal test, indicating improved fine motor coordination, and significantly less pronounced hindlimb clasping, reflecting reduced striatal dysfunction^45^ (**Fig. 1C, Supplementary Fig. 3A**). We did not observe effects in the beam traversal, pole descent, or wire hang tests (**Supplementary Fig. 1A, Supplementary Fig. 3A**). GI symptoms also improved; namely, benCom-PD-treated mice expelled a rectally-inserted glass bead more rapidly than controls and produced fecal pellets with higher scores on the Bristol stool scale, suggesting alleviation of constipation-like symptoms (**Fig. 1C**, **Supplementary Fig. 1B**). Accordingly, motor and GI function improvements in mice treated with benCom-PD were accompanied by a reduction in levels of phosphorylated-S129 α-synuclein (pS129-αSyn), a pathogenic modification that correlates with disease measures^46^, in the striatum (**Fig. 1D, E**), but not in the substantia nigra (**Supplementary Fig. 1C, D**); no differences were observed in abundance of aggregated αSyn (**Supplementary Fig. 1E-H**). Together, these findings reveal that supplementation of Thy1-ASO mice with specific commensal microbes, commonly depleted in human PD, improves symptoms and pathology associated with PD.

**Figure 1.**
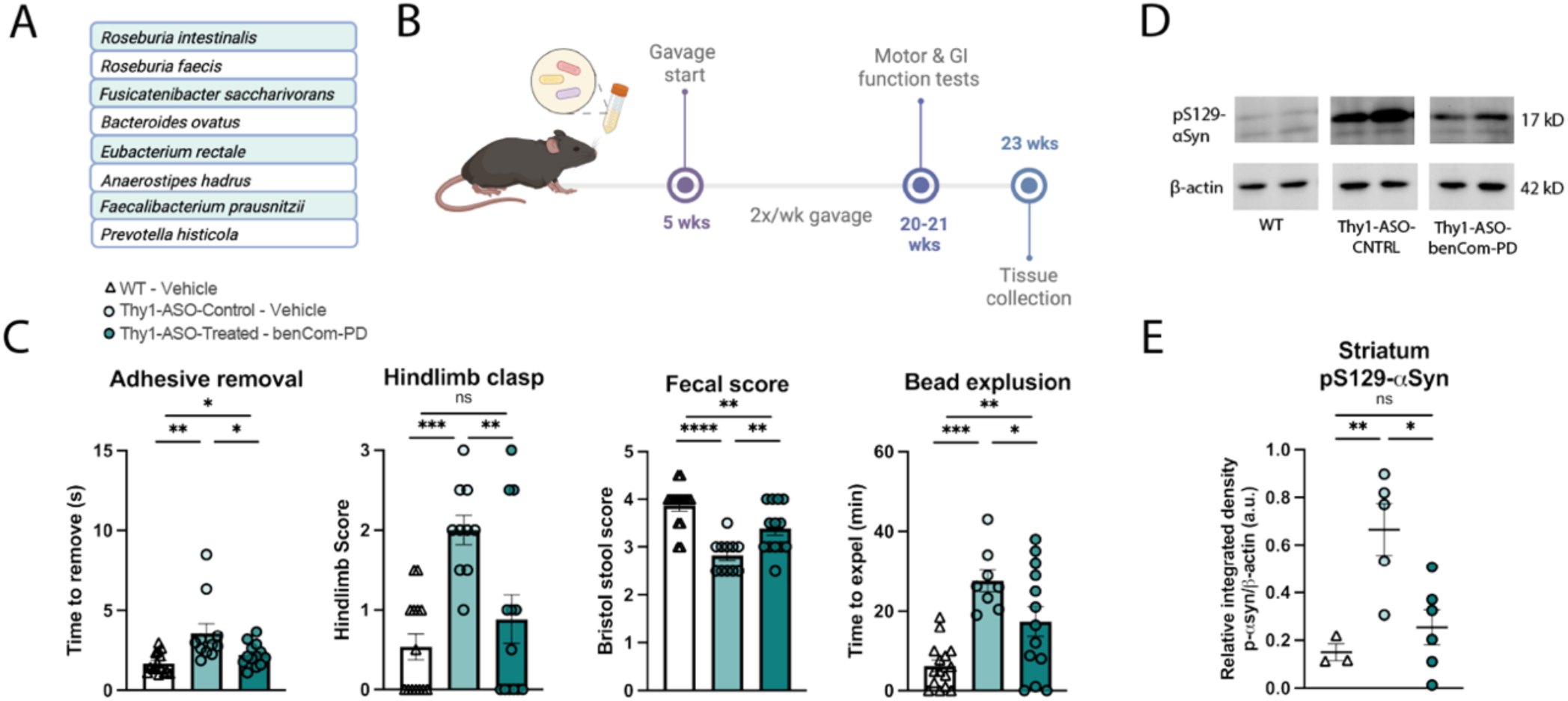
Treatment with benCom-PD improves motor and GI function and reduces pS129-αSyn in the striatum. (**A**) Composition of the benCom-PD consortium consisting of 8 taxa that are reduced in the PD microbiome. (**B**) Animal treatment and testing timeline. (**C**) Improvements in motor and GI function assays with benCom-PD treatment of Thy1-ASO mice compared to controls, tested at 20 weeks. N = 11-15. Points represent individual animals; bars represent standard error of the mean. (**D, E**) Representative images (D) and quantification (E) of Western blots for phosphorylated-S129-α-synuclein in striatal tissue. N = 3-6. Points represent individual animals; bars represent standard error of the mean. Behavior data in (C) analyzed by Kruskal-Wallis test followed by the Conover-Iman post-hoc test with Benjamini-Hochberg false discovery rate (FDR) correction; protein data in (E) analyzed by one-way ANOVA with Tukey post-hoc test. *p≤0.05; **p≤0.01; ***p≤0.001; ****p≤0.0001. Abbreviations: WT – wildtype; Thy1-ASO-CNTRL – Thy1-human α-synuclein overexpressing control; benCom – beneficial commensal consortium; pS129-αSyn – phosphorylated-S129-alpha-synuclein

### F. prausnitzii has robust therapeutic effects on motor symptoms and synucleinopathy

Across published studies of PD patients curated in the BugSigDB database^47^, *Faecalibacterium prausnitzii* emerges as the most consistently reduced microbial taxon, reported as decreased in 16 independent studies and never as increased (**Supplementary Table 1**). Additionally, *F. prausnitzii* has well-documented immunomodulatory properties and is effective in ameliorating experimental colitis^48^. We therefore identified it as a particularly compelling candidate for further investigation among the taxa represented in benCom-PD. We treated animals with *F. prausnitzii* mono-cultures for 15 weeks, using the same dosing regimen established for benCom-PD (**Fig. 2A**) and assessed motor and GI function. *F. prausnitzii*-treated mice showed significant improvements in the adhesive removal and hindlimb clasping tests compared to vehicle-treated Thy1-ASO mice (**Fig. 2B, Supplementary Fig. 3B**), similar to the effects observed with benCom-PD. Animals treated with *F. prausnitzii* alone further exhibited an improved ability to cross a challenging beam, a test of motor coordination, as reflected by reduced crossing time and fewer errors per step (paw slips) (**Fig. 2B, Supplementary Figs. 2A, 3B, & 4**). GI function was also improved following *F. prausnitzii* treatment, marked by decreased time required to expel a rectally-inserted glass bead and improved Bristol stool scale scores following treatment (**Fig. 2B, Supplementary Figs. 2B & 4**). While we did not detect changes in levels of pS129-αSyn in brain tissues (**Supplementary Fig. 2C-F**), supplementation with *F. prausnitzii* reduced aggregation of αSyn in the substantia nigra (**Fig. 2C, D**), but not in the striatum (**Supplementary Fig. 2G, H**), of Thy1-ASO mice relative to vehicle-control Thy1-ASO mice. These findings demonstrate that treatment with *F. prausnitzii,* a single microbe found to be depleted in the PD patient gut microbiome, is sufficient to markedly improve disease-associated outcomes in the Thy1-ASO mouse model.

**Figure 2.**
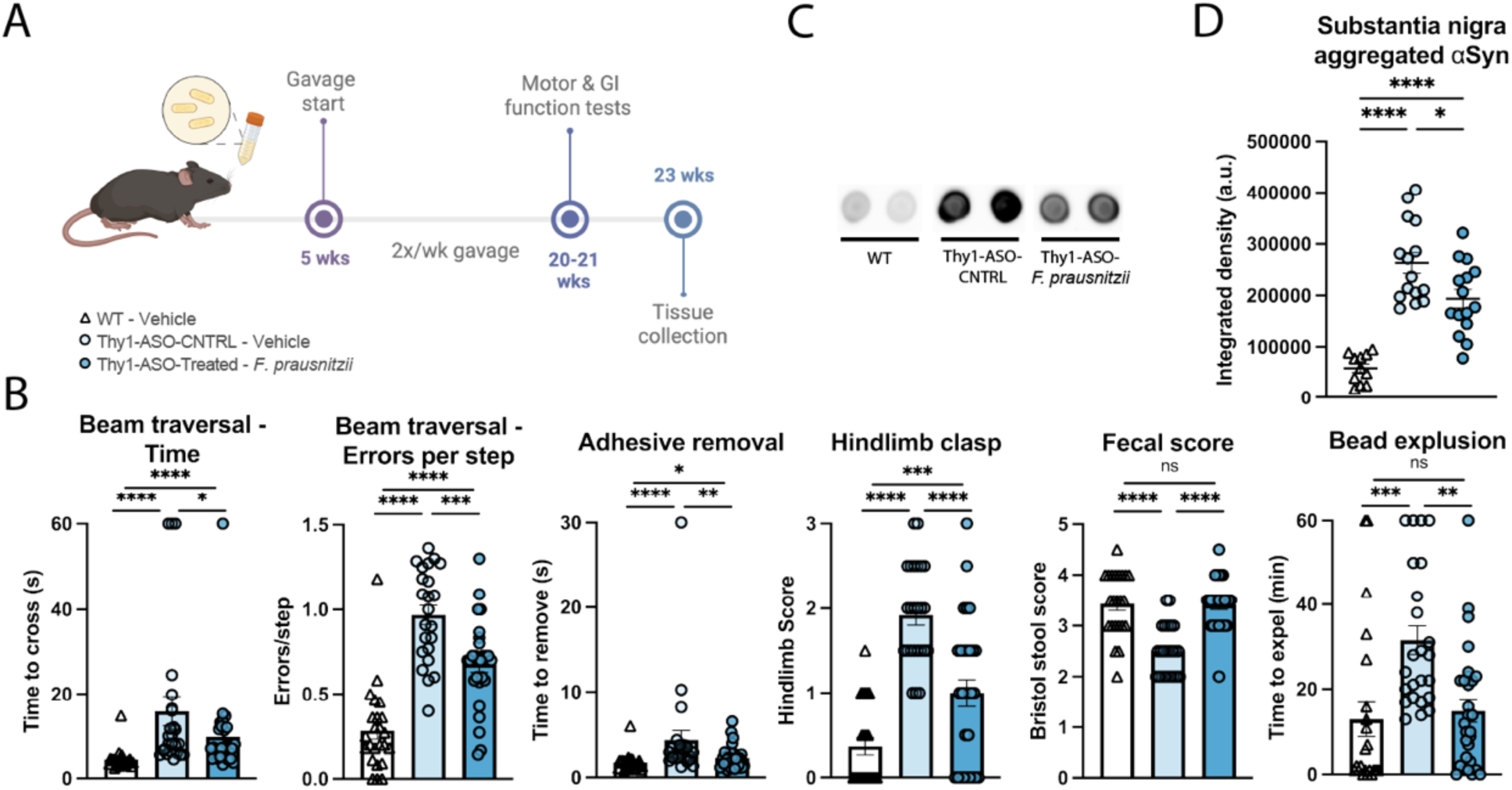
Treatment with *F. prausnitzii* mono-culture ameliorates motor and GI symptoms and reduces αSyn aggregates in the substantia nigra. (**A**) Animal treatment and testing timeline. (**B**) Improvements in motor and GI function assays with *F. prausnitzii* treatment in Thy1-ASO animals compared to controls, tested at 20 weeks. N = 23-29. Points represent individual animals, compiled from two independent cohorts; bars represent standard error of the mean. (**C, D**) Representative images (C) and quantification (D) of dot blots for aggregated α-synuclein in the substantia nigra. N = 11-15. Points represent individual animals, compiled from two independent cohorts; bars represent standard error of the mean. Behavior data in (B) analyzed by Kruskal-Wallis test followed by the Conover-Iman post-hoc test with Benjamini-Hochberg false discovery rate (FDR) correction; protein data in (D) analyzed by one-way ANOVA with Tukey post-hoc test. *p≤0.05; **p≤0.01; ***p≤0.001; ****p≤0.0001. Abbreviations: WT – wildtype; Thy1-ASO-CNTRL – Thy1-human α-synuclein overexpressing control; *F. prausnitzii* – *Faecalibacterium prausnitzii;* αSyn – α-synuclein

### F. prausnitzii treatment aligns the Thy1-ASO microbiome to more closely resemble a WT profile

The fecal microbiome is altered in Thy1-ASO mice and other PD models compared to WT mice^49–53^. Metagenomic analysis revealed that *F. prausnitzii* treatment remodeled or maintained the gut microbiome of Thy1-ASO mice to more closely resemble that of WT animals, while the composition in untreated Thy1-ASO mice deviated from the other two groups (**Fig. 3, Supplementary Fig. 5**). At the phylum level, relative abundance profiles appeared largely stable across groups, with no major shifts observed between conditions (**Fig. 3A**). However, at the species genome bin (SGB) level, Principal Coordinate Analysis (PCoA) of Bray-Curtis dissimilarity revealed a separation of the microbiomes of WT and vehicle-treated control Thy1-ASO animals (**Fig. 3B**), reflecting genotype-associated differences that align with previous findings^50,51^. Notably, supplementation with *F. prausnitzii* partially prevented compositional changes in the Thy1-ASO mice (**Fig. 3B**). Pairwise comparisons of Bray-Curtis dissimilarity showed significant differences between WT and control Thy1-ASO mice (pairwise PERMANOVA FDR = 0.011), as well as between *F. prausnitzii*-treated Thy1-ASO and control Thy1-ASO animals (pairwise PERMANOVA FDR = 0.020), while microbial compositions between WT and treated Thy1-ASO mice showed less dissimilarity (pairwise PERMANOVA FDR = 0.250) (**Fig. 3C**). However, measures of observed richness, Shannon diversity, and the Firmicutes/Bacteroidetes ratio did not differ significantly between groups (**Supplementary Fig. 5**). Differential abundance and prevalence analyses identified subtle shifts in multiple taxa, suggesting that a community-wide alteration may underlie the Thy1-ASO microbiome’s increased similarity to the WT profile, rather than pronounced alterations in specific taxa (**Fig. 3D**). Interestingly, presence of *F. prausnitzii* was not identified in the metagenomes, likely due to its weak ability to individually colonize the mouse gut^54–56^.

**Figure 3.**
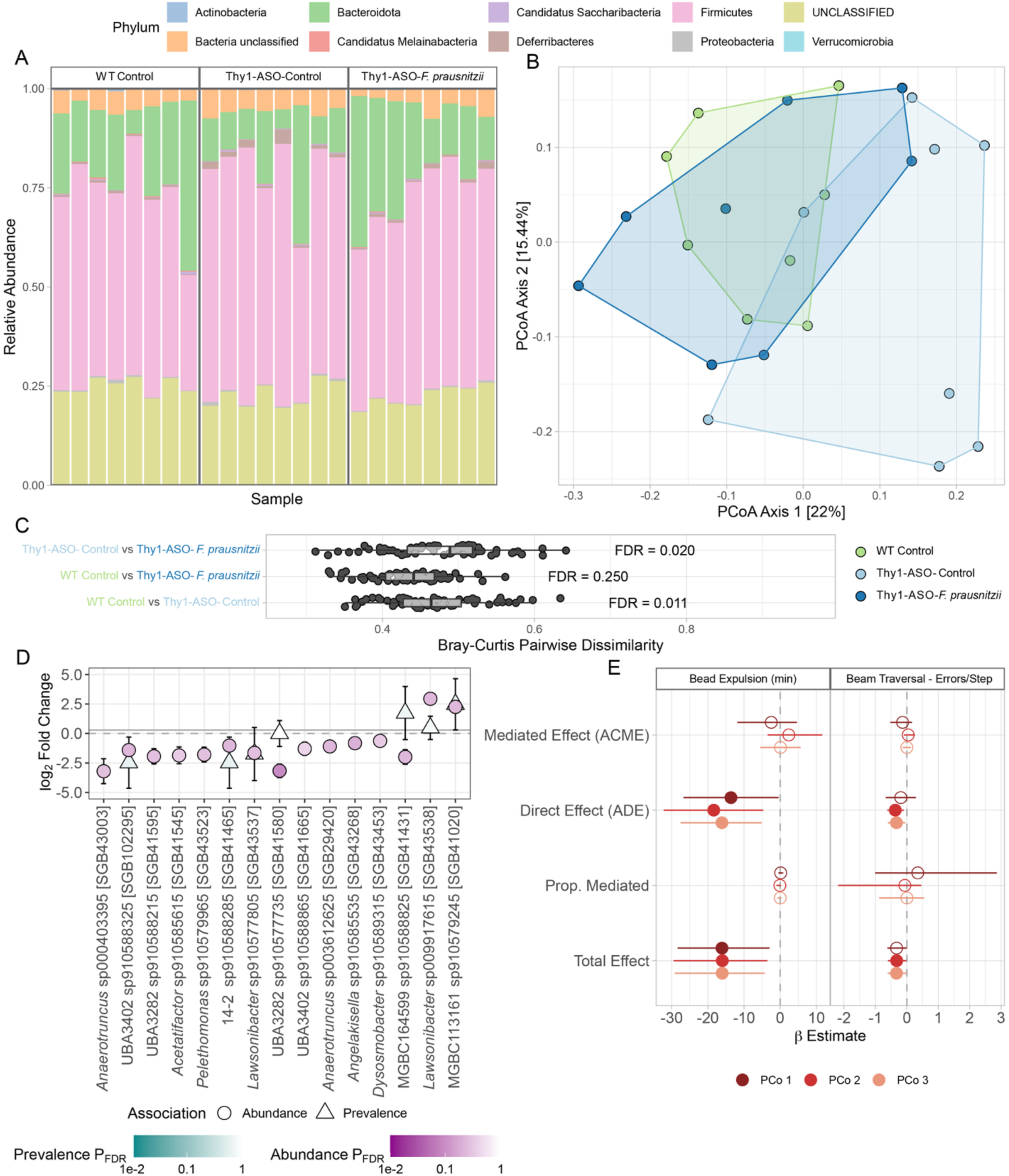
The gut microbiome of Thy1-ASO mice is mildly remodeled upon treatment with *F. prausnitzii*. (**A**) Stacked bar plot of relative abundance of bacterial phyla in the gut microbiome, including unclassified reads. Each bar represents a single mouse. (**B**) Principal Coordinate Analysis (PCoA) of the Bray-Curtis dissimilarity of fecal microbiome composition of each group. Percentages in brackets represent variance explained by each axis. (**C**) Boxplot of pairwise dissimilarity. Statistical significance was calculated using pairwise permutational multivariate analysis of variance (PERMANOVA) with Benjamini-Hochberg adjusted false discovery rate (FDR) across multiple pairwise comparisons. (**D**) Differential abundance and/or prevalence of the top 15 species-level genome bins (SGBs) in *F. prausnitzii*-treated Thy1-ASO mice compared to vehicle-treated Thy1-ASO controls. Analysis was performed in MaAsLin3 on log2-transformed relative abundance values scaled between 0 and 1 with no further adjustments. SGB nomenclature was enriched using the Genome Taxonomy Database provided in MetaPhlAn v4.1.1 See **Supplementary Data 4 and 5** for full taxonomy. (**E**) Mediation analysis of the first three microbiome principal coordinates as mediators of the effect of *F. prausnitzii* treatment on the beam traversal errors/step and bead expulsion assays. Effect sizes (β) and 95% confidence intervals are shown. Full circles indicate a p-value ≤0.05, empty circles indicate p>0.05. Refer to **Supplementary Data 6 and 7** for full mediation analysis data. Abbreviations: WT – wildtype; Thy1-ASO – Thy1-human α-synuclein overexpressing; *F. prausnitzii* – *Faecalibacterium prausnitzii;* ACME – Average Causal Mediation Effect; ADE – Average Direct Effect; Prop. – Proportion

To determine whether the microbiome changes we observed play a role in the motor and GI symptom improvements seen with treatment, we conducted mediation analysis with *F. prausnitzii* treatment as the predictor and altered microbiome composition as a potential mediator (**Fig. 3E**). As functional outcomes, we selected errors/step in beam traversal and time to bead expulsion, as these were the most robust and disease-relevant effects among the motor and GI function assays. Mediation analysis revealed that *F. prausnitzii* administration had a significant direct influence on both behavioral outcomes, with an insignificant indirect effect, reflected by an average causal mediation effect (ACME) close to zero. These findings suggest that the physiological outcomes were not mediated by overall microbiome alterations, as approximated by the first three principal coordinates of Bray-Curtis dissimilarity, but rather influenced directly by *F. prausnitzii*. Together, our results demonstrate that *F. prausnitzii* is sufficient to drive behavioral improvements and contributes to a mild remodeling or maintenance of the gut microbiome profile in Thy1-ASO mice to more closely resemble that of WT animals.

### F. prausnitzii alters the immune landscape in treated animals

Probiotic treatment with *F. prausnitzii* suppresses intestinal and systemic inflammation in mouse models of colitis, arthritis, and during testing of anti-cancer drugs^48,57–59^. Here, flow cytometric analysis of Thy1-ASO mice receiving *F. prausnitzii* revealed that treatment led to a significant increase in CD4+CD25+FoxP3+ T_REG_ cell numbers in the mesenteric lymph nodes (MLNs) (**Fig. 4A, B and Supplementary Figs. 6 & 7**). T_REG_ deficiencies have been implicated in PD and its associated mouse models^60–62^. T_REG_ expansion was accompanied by elevated levels of the anti-inflammatory cytokine IL-10 in proximal large intestinal tissue, as measured by multiplex bead assay (**Fig. 4C**). These findings demonstrate that *F. prausnitzii* exerts immunomodulatory effects on adaptive immune cell populations in Thy1-ASO mice.

**Figure 4.**
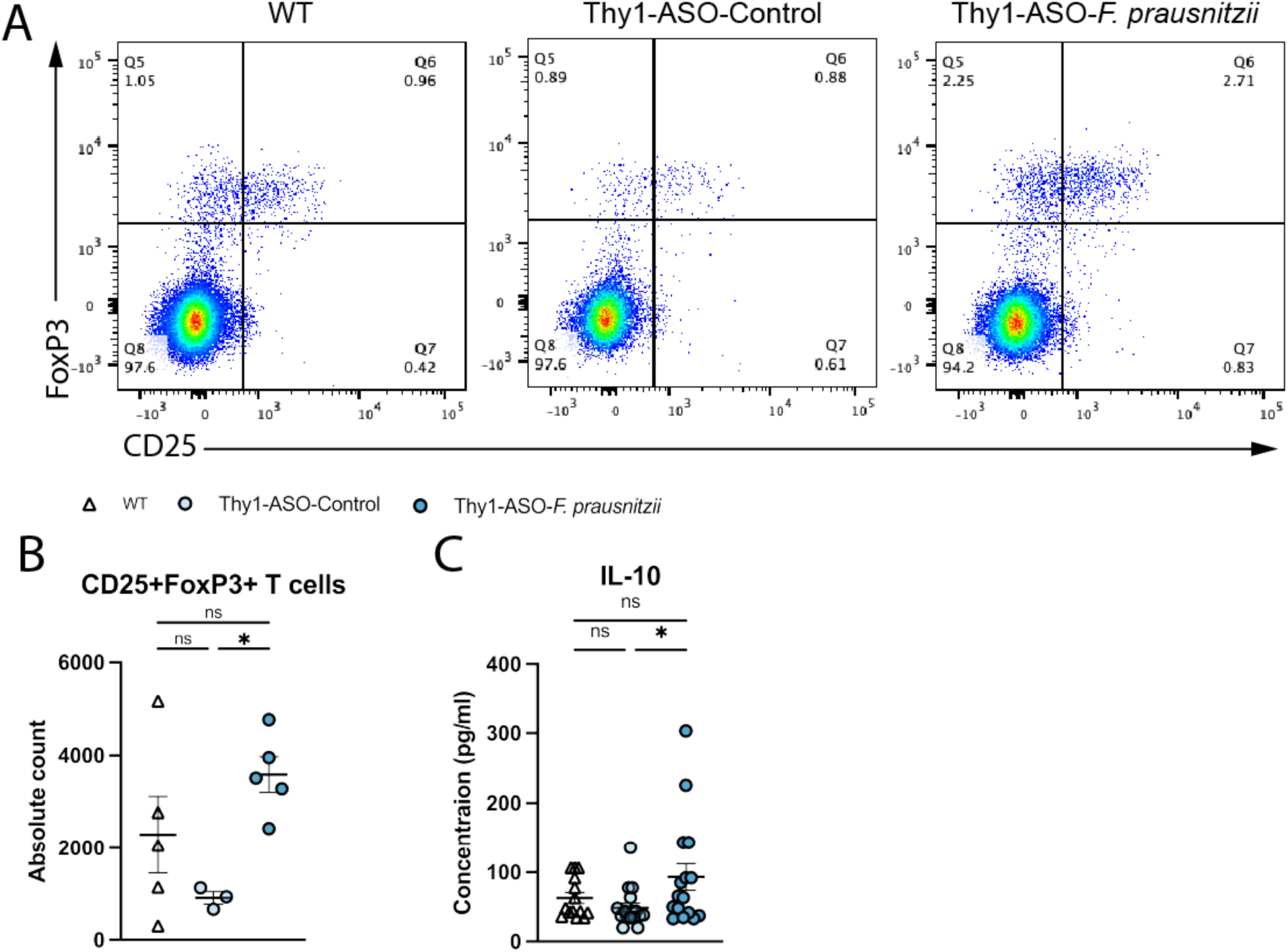
*F. prausnitzii* supplementation induces anti-inflammatory processes in gut immune compartments. (**A, B**) Representative flow cytometry plots (A) and quantification (B) of activated CD25+FoxP3+ regulatory T cells in mesenteric lymph nodes. N = 3-5. (**C**) IL-10 concentration in colon tissue as measured by multiplex bead assay. Data are compiled from two independent cohorts. N= 14-17. Points represent individual animals, bars represent standard error of the mean. Data analyzed by one-way ANOVA with Tukey post-hoc test. ns – not significant; *p≤0.05 Abbreviations: WT – wildtype; Thy1-ASO – Thy1-human α-synuclein overexpressing; *F. prausnitzii* – *Faecalibacterium prausnitzii*; IL-10 – interleukin-10

Despite the alterations in the immune profile and shifts in the gut microbiome (see **Fig. 3**) observed in *F. prausnitzii*-treated animals, we did not detect differences in the levels of volatile fatty acids, including measurable SCFAs, in fecal, cecal, or serum samples from Thy1-ASO-treated mice (**Supplementary Fig. 8A-C**). Fermentation pathway analysis of metagenomic data confirmed no changes in abundance of genes associated with SCFA metabolite production pathways across experimental conditions (**Supplementary Fig. 8D**). This outcome is in contrast to previous studies reporting increases in SCFAs following *F. prausnitzii* supplementation^57,63,64^, though at least one other study similarly found no effect^65^. While the therapeutic effects of *F. prausnitzii* in Thy1-ASO mice do not appear to be mediated by SCFAs, further mechanistic studies with knockouts of bacterial (butyrate production) or mouse (SCFA receptors) genes are needed.

### F. prausnitzii supplementation remodels the gene expression profile of the large intestine

To gain insight into potential molecular mechanisms of *F. prausnitzii*-mediated protection, we performed bulk RNA sequencing of the proximal large intestine, which revealed significant (p = 2.00E-04) gene expression differences between experimental groups (**Fig. 5A**). Compared to control (vehicle-treated) Thy1-ASO mice, *F. prausnitzii* supplementation altered expression of several immune-related genes linked to PD (**Fig 5B, E**). The gene encoding transferrin receptor (*Tfrc*), whose transcription was upregulated in colon tissue from *F. prausnitzii*-treated animals, is essential for iron uptake and supports T and B cell proliferation^66^. TFRC and transferrin (TF) are reduced in the temporal cortex and substantia nigra of PD patients, and TF injections ameliorate 1-methyl-4-phenyl-1,2,3,6-tetrahydropyridine (MPTP) mouse model outcomes^67–69^. CCL5, or RANTES, is a chemokine that is elevated in the serum of PD patients^70,71^, and its gene (*Ccl5*) was downregulated in mice treated with *F. prausnitzii*. *Cd8a,* which encodes a protein expressed on cytotoxic T cells, was also downregulated following *F. prausnitzii* supplementation. CD8+ cytotoxic T cells have been found to infiltrate the brain in PD models and are elevated in the peripheral blood of patients^53,72,73^. Thus, in the gut tissue of *F. prausnitzii*-treated Thy1-ASO mice, we observed normalized expression of several genes (or their products) previously reported to be altered in the blood and brain of PD patients.

**Figure 5.**
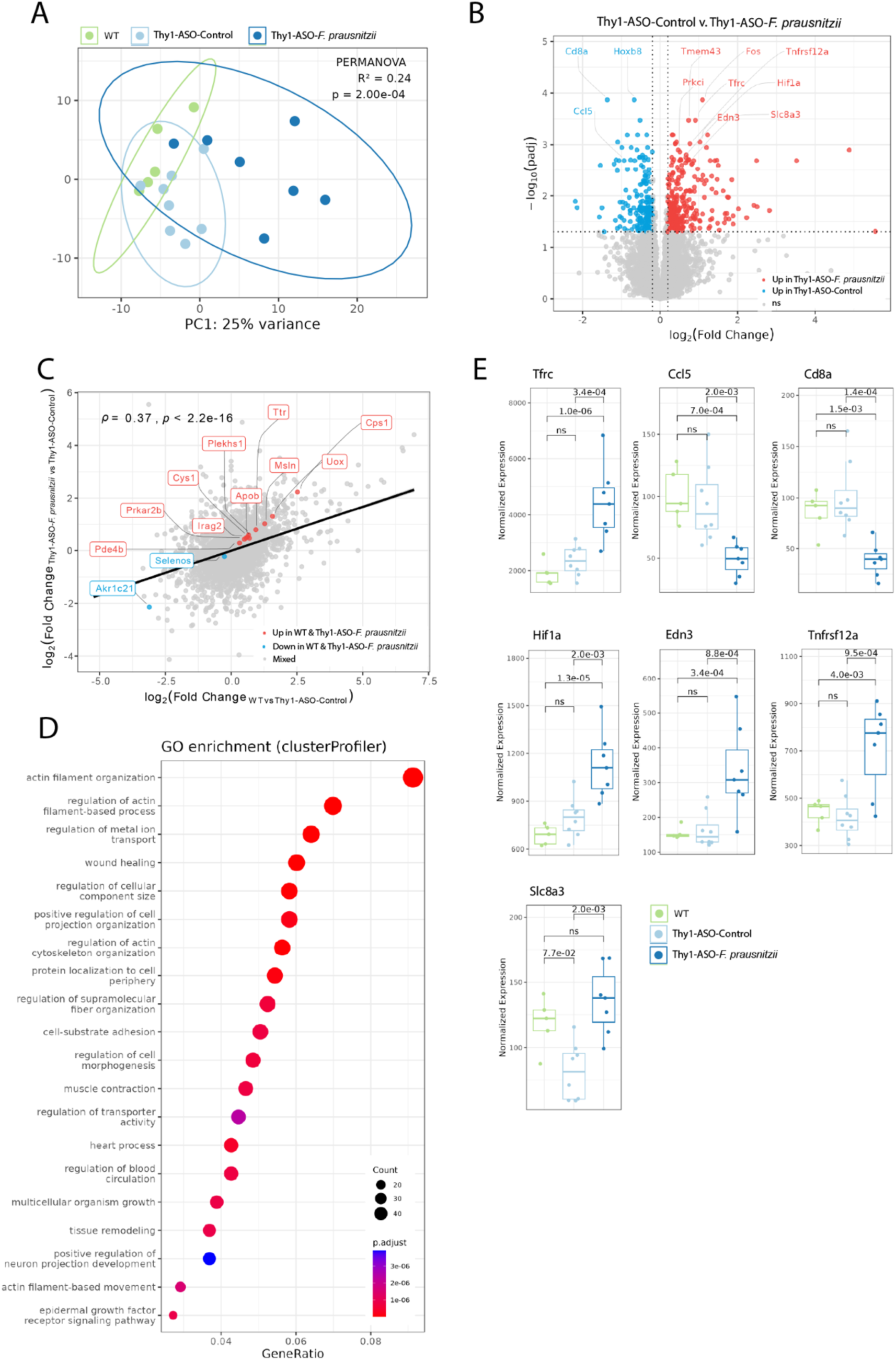
Host gene expression profiles in the large intestine are modulated by *F. prausnitzii* supplementation. (**A**) Principal component analysis of gene expression profiles in large intestine. Ellipses indicate 95% confidence intervals. R^2^ and p-value PERMANOVA statistics are displayed for the experimental condition category using 9,999 permutations. (**B**) Volcano plot of genes differentially expressed between Thy1-ASO-Control and Thy1-ASO-*F. prausnitzii* groups. (C) Cross-condition scatter plot of genes differentially expressed between WT and Thy1-ASO-Control on the x-axis and Thy1-ASO-*F. prausnitzii* and Thy1-ASO-Control on the y-axis. Pearson correlation coefficient *(Ρ)* and p-value of log2 fold-change of all genes tested are shown. (D) Gene ontology (GO) enrichment analysis showing pathways upregulated in Thy1-ASO-*F. prausnitzii* tissue v. Thy1-ASO-Control. (**E**) Expression levels of differentially regulated genes of interest. Box plots show normalized gene expression values (size-factor scaled counts) with DESeq2 adjusted p-values between comparisons. ns – not significant Abbreviations: WT – wildtype; Thy1-ASO – Thy1-human α-synuclein overexpressing; *F. prausnitzii* – *Faecalibacterium prausnitzii*

Comparative analysis of differentially expressed genes revealed a strong positive correlation between differences observed in WT versus Thy1-ASO control mice and those seen in Thy1-ASO *F. prausnitzii*-treated versus Thy1-ASO control mice (ρ = 0.37, p < 2.2E−16) (**Fig. 5C**), suggesting that probiotic treatment shifts the transcriptional profile of Thy1-ASO animals towards that of WT mice. Of note, among the genes that were upregulated in colonic tissue of both WT and Thy1-ASO *F. prausnitzii* mice compared to Thy1-ASO control animals, was transthyretin (*Ttr*), a gene relevant to PD. *Ttr* encodes a transporter of thyroid hormones and retinol, and mutations in this gene cause transthyretin amyloidosis, which is characterized by amyloid deposits^74^. TTR is reduced in the CSF of Lewy body disease and Alzheimer’s disease patients^75^, and is able to cleave free αSyn *in vitro*, suggesting a possible neuroprotective role^76^. Selenoprotein S, or SELENOS, supports protein folding, limits oxidative stress, and localizes to Lewy bodies in PD^77^. SELENOS activity is dependent on selenium, a trace element whose deficiency has been linked to cognitive decline, and selenium supplementation has shown therapeutic effects in several PD models^77,78^. *Selenos* gene expression was slightly downregulated in both WT and *F. prausnitzii*-treated Thy1-ASO mice, potentially indicating a reduced need for its activity in these conditions.

Gene ontology (GO) analysis indicated significant enrichment in pathways associated with tissue repair and cytoskeletal remodeling in *F. prausnitzii*-treated Thy1-ASO animals compared to Thy1-ASO control mice (**Fig. 5D, E**). Enrichment of *Actin filament-*related, *wound healing*, and *muscle contraction* pathways suggests activation of cellular mechanisms involved in restoring intestinal barrier integrity and repairing gut function. Hypoxia inducible factor-1α (*Hif1a*), which is part of the *wound healing* and *tissue remodeling* pathways, encodes a transcription factor that regulates cellular response to low oxygen and is also an inducer of FoxP3+ regulatory T cells in inflamed intestinal tissue^79^. Additionally, HIF-1α has been shown to be involved in mechanisms related to several drug treatments in PD animal models and has been proposed as a potential therapeutic target^80–82^. Endothelin-3 (*Edn3*), involved in the *regulation of metal ion transport* and *muscle contraction* pathways, promotes T_REG_ proliferation *in vitro*^83^. Additionally, GO pathways containing TNFα-related genes were impacted by *F. prausnitzii* treatment. While classically thought of as a pro-inflammatory cytokine, TNFα can be required for T_REG_ function^84,85^ and is involved in tissue repair^86,87^. Tumor necrosis factor superfamily member 12a (*Tnfrsf12a*), upregulated in the *wound healing* pathway, encodes a receptor of TNFα-like weak inducer of apoptosis (TWEAK/Tnfrsf12), which plays a role in tissue regeneration^88,89^. *Slc8A3* (NAX3) encodes a sodium/calcium exchanger protein that regulates TNFα release from macrophages and monocytes^90^, and was upregulated in the *muscle contraction* pathway by *F. prausnitzii*. Interestingly, calcium homeostasis disruption is believed to contribute to PD pathology, with Na^+^/Ca^2+^ exchanger proteins implicated in this dysregulation^91^. The upregulation of genes involved in immune homeostasis and tissue repair suggest that *F. prausnitzii* has therapeutic effects in the intestine of Thy1-ASO animals, though potential gut-to-brain pathways remain to be explored. Collectively, our findings highlight the therapeutic potential of *F. prausnitzii* in a preclinical model of PD, demonstrating its capacity to ameliorate both behavioral and neuropathological features, while also modulating gut microbiota composition, immune responses, and host gene expression in the Thy1-ASO mouse model.

## Discussion

Fewer than 5% of PD cases are caused by monogenic risk, and only about 20% can be attributed to any genetic factor, suggesting that most PD likely arises from environmental exposures or gene–environment interactions. Indeed, pesticides, heavy metals, solvents, air pollution, and other environmental toxicants have been implicated in PD^92^. Concordant studies have uncovered differences in gut microbiome composition between PD patients and household or population controls^32,36,93,94^, representing a potential environmental risk. Indeed, the microbiome has profound effects on disease outcomes in several animal models of PD. For example, we and others have demonstrated that pathogenic gut bacterial species can accelerate motor symptoms and brain pathology in PD mouse models^95–97^. A high-fiber diet corrects microbiome alterations in Thy1-ASO mice and prophylactically improves motor deficits via modulation of microglial activation^51^. Furthermore, fecal microbiome transplants (FMT) from human PD donors into Thy1-ASO mice worsen motor performance compared to FMT from healthy donors^95^. In addition to animal studies^98–101^, several clinical trials have assessed safety and efficacy of FMTs in PD, yielding encouraging but inconsistent results that suggest further optimization of treatment strategies may be needed^102–105^.

Here, we investigated, for the first time to our knowledge, whether the reintroduction of microbes depleted in PD could restore physiological balance. Oral administration of a defined microbial consortium composed of taxa consistently depleted in the gut microbiome of multiple PD cohorts improved select features of disease in the Thy1-ASO mouse model. Further, a single member of this consortium, *F. prausnitzii*, was sufficient to ameliorate both motor and GI function deficits and reduce αSyn aggregation in the brain. *F. prausnitzii* treatment shifted the microbiome in Thy1-ASO mice closer to a WT profile, exerted immunomodulatory effects, and enhanced intestinal transcriptional programs associated with tissue regeneration. Our findings demonstrate that targeted microbial supplementation can elicit functional improvements in preclinical models, and may translate to replenishing microbes that are depleted in PD patients.

Many of the strains we tested in benCom-PD, beyond *F. prausnitzii*, also have therapeutic or anti-inflammatory effects in other contexts*. Roseburia intestinalis* has been shown to inhibit pro-inflammatory interleukin-17 (IL-17) production, promote T_REG_ expansion, and ameliorate colitis in mice^106–108^. A related but less studied bacterium, *Roseburia faecis,* improves symptoms related to irritable bowel syndrome (IBS) in a stress-induced rat model^109^. *Fusicatenibacter saccharivorans* and *Anaerostipes hadrus* are both depleted in patients with colorectal cancer and IBD^110–113^, while *Bacteroides ovatus* reduces colitis severity in animals through modulation of immunity^114,115^ and improves immunotherapy response in graft versus host disease models^116^. Treatment with *Eubacterium rectale* has also been shown to improve response to anti-PD1 therapy^117^, and mitigates intestinal lymphoma in tumor-bearing mice^118^. *Prevotella histicola* promotes T_REG_ function, and suppresses arthritis^119^ and multiple sclerosis in animal models^120^.

Metagenomic sequencing revealed that treatment with *F. prausnitzii* modestly, but significantly, remodeled or maintained the fecal microbiome of Thy1-ASO mice, with treatment-associated shifts in overall community structure. Measures of alpha diversity remained largely unchanged across groups, consistent with a selective, rather than global, change in the microbial community by *F. prausnitzii*. Differential abundance analysis also indicated subtle directed ecological interactions with endogenous microbial populations. Importantly, and consistent with the finding that the effects of *F. prausnitzii* on microbiome composition appear targeted, mediation analysis suggested that the therapeutic effects of *F. prausnitzii* are likely to be largely direct rather than mediated by remodeling of the broader microbiome.

Following treatment with *F. prausnitzii*, we observed increased intestinal T_REG_ populations and elevated levels of the anti-inflammatory cytokine IL-10, consistent with prior findings of its capacity to induce immune tolerance^40,121,122^. While butyrate is a major SCFA produced by *F. prausnitzii* and has been implicated in T_REG_ development^123^, we did not detect increased SCFA levels following treatment, indicating that the bacterium’s therapeutic effects in Thy1-ASO mice may be independent of SCFAs and perhaps driven by secreted immunomodulatory molecules such as the Microbial Anti-inflammatory Molecule (MAM)^38^.

Transcriptomic profiling of colonic tissue revealed that *F. prausnitzii* shifted the gene expression landscape of PD mice toward that of WT control animals, with enrichment of genes and pathways involved in immune regulation, epithelial repair and regeneration. Thus, *F. prausnitzii* alters the gene expression profile in this PD mouse model, potentially contributing to its immunomodulatory and therapeutic effects, a hypothesis that remains to be tested.

In prior probiotic treatment studies using PD animal models, *Lacticaseibacillus rhamnosus* has shown neuroprotective effects in MPTP-treated mice^124,125^. Similarly, *Lactobacillus plantarum* PS128 reduces microglial activation and motor deficits in MPTP-treated^126^ and rotenone-exposed mice^127^. Another study reported the ability of *Akkermansia muciniphila* to ameliorate PD-like symptoms in the MPTP model^128^. In early clinical work, multi-strain probiotics containing *Lactobacillus acidophilus*, *Lactobacillus casei*, *Bifidobacterium bifidum*, and *Bifidobacterium lactis* have demonstrated improvements in constipation^30,31^. *L. plantarum* PS128 elicits modest improvements in UPDRS scores, likely mediated by increased plasma dopamine metabolites^129^. However, meta-analyses emphasize that while commercial probiotics modestly improve GI and motor outcomes, the quality of evidence remains low due to small sample sizes, short treatment durations, and lack of mechanistic biomarkers^130^. Further, unlike *F. prausnitzii*, these species have not been shown to be reproducibly reduced in PD patients.

There are several limitations to this study. First, we only tested benCom-PD or *F. prausnitzii* in a single preclinical model – the Thy1-ASO mouse – which captures several key aspects of PD, including αSyn accumulation and progressive behavioral deficits^43^, but lacks others, such as overt dopaminergic neuron loss at the ages tested. Additionally, integration of the αSyn transgene on the X chromosome makes expression inconsistent in female mice due to X inactivation, so only male mice were studied. Future studies employing complementary models and including aging (the primary risk factor for PD) will help validate and extend the observations of the current work. Second, although *F. prausnitzii* improved PD-associated behaviors and corrected certain molecular changes caused by αSyn overexpression, the precise mechanisms underlying its effects remain untested. Whether these improvements are mediated primarily through peripheral immune modulation, direct gut-brain signaling (i.e., vagal or spinal neurons), production of beneficial microbial metabolites, microbiome restoration, or a combination thereof requires further investigation. Finally, our treatment strategy tested a prophylactic approach prior to disease symptom onset; thus, translation to human studies would require identification of at-risk or prodromal PD populations, which the PD research community is advancing, but is not yet reliably achievable.

Our study provides proof-of-concept that targeted restoration of bacterial species that are depleted in PD patients can improve disease-relevant outcomes, supporting the emerging hypothesis that the gut microbiome is a modifiable contributor to PD. Addressing environmental risk factors of PD, such as the gut microbiome, compared to correcting genetic predispositions, offers a likely more tractable therapeutic route. Furthermore, development of PD-specific, rather than generic, probiotics represents a promising avenue for advancing treatment options.

## Methods

### Bacterial cultures

Strains were purchased from DSMZ and ATCC and grown and prepared inside an anaerobic chamber (Coy Laboratories) (80% nitrogen, 10% carbon dioxide, and 10% hydrogen) at 37℃. *Roseburia intestinalis* (DSMZ14610), *Roseburia faecis* (DSMZ16840), and *Anaerostipes hadrus* (ATCC29173) were grown in Yeast Casitone Fatty Acids with Carbohydrates (YCFAC) media (Anaerobe Systems). *Fusicatenibacter saccharivorans* (DSMZ26063) and *Prevotella histicola* (DSMZ19854) were grown in Peptone Yeast Extract Broth with Glucose (PYG) media (Anaerobe Systems). *Bacteroides ovatus* (ATCC8483) was grown in Chopped Meat Broth (Anaerobe Systems) and *Eubacterium rectale* (ATCC33656) was grown in Mega Medium^131,132^. *Faecalibacterium prausnitzii* (DSMZ17677) was grown in Chopped Meat with Carbohydrates (CMC) (Anaerobe Systems). For preparing aliquots for intragastric administration (gavage), cultures were grown to an approximate optical density (600nm) of 1, then spun down and resuspended in sterile 50% glycerol to an approximate Colony-Forming Unit (CFU) density of 10^8^/mL. For benCom-PD, all bacterial cultures were pooled in equal amounts. Aliquots were frozen at -20℃ for short-term storage in cryotubes wrapped with parafilm to protect from oxygen exposure. On the day of gavage, an aliquot was thawed and resuspended in a sterile solution of phosphate buffered saline (PBS) with 1.5% NaHCO_3_. Gavage tubes were set up inside the anaerobic chamber to reduce oxygen exposure and placed into a GasPak box (BD) with an AnaeroPack (MGC) for transportation to the animal facility. Two times a week, animals received a gavage of 100μL of vehicle (PBS + 1.5% NaHCO_3_) (WT and Thy1-ASO-Control groups) or bacterial culture (benCom-PD or *F. prausnitzii* - Thy1-ASO-Treated groups).

### Mice

All animal husbandry and experiments were approved by the Caltech Institutional Animal Care and Use Committee. Alpha-synuclein overexpressing (Thy1-ASO) male mice were bred by crossing wildtype (WT) BDF1 males with Thy1-ASO females of the BDF1 background that were heterozygous for the Thy1 promoter driving the human alpha-synuclein transgene located on the X chromosome^43,133^. Mice had a conventional, specific pathogen-free (SPF) microbiome and were housed in microisolator cages with HEPA-filtered ventilation. All animals received irradiated chow (LabDiet 5053) and water *ad libitum*, and were maintained on a 12 hour light-dark cycle.

### Motor and gastrointestinal function testing

Motor function test methods were similar to those previously described^95,134^. All motor and GI function tests were performed between hours 7 and 9 of the light phase, at the same time every day, inside a biosafety cabinet in the same facility. Animals were habituated to the behavior testing room for 1 hour prior to the start of motor and GI function experiments. Beam traversal and pole descent training and testing were done together on two consecutive days, with approximately one hour of rest time between each task. After one rest day, testing for wire hang was conducted. Lastly, adhesive removal and hindlimb clasping scoring were performed on the final day. GI function tests were performed the following week, with bead expulsion assayed last.

For the beam traversal assay, four 0.25m plexiglass beam sections, starting at 3.5cm width and narrowing to 0.5cm width, were placed together to form a 1m beam. The beam was supported by 3 standard size mouse cages at the beginning, middle, and end. Animals were placed onto the widest section of the beam and trained to run across the beam towards their home cage, which was placed sideways at the end of the narrowest section, with the open side facing the animal.

Mice were given 3 trials on the training day. The first trial was guided, with the home cage being brought close and moved along with the animal to encourage forward progress. The second trial was performed similarly but with less guidance and encouragement. For the third trial, the home cage was returned to the end of the beam and animals traversed the beam with limited assistance. On the next day, animals were tested for their ability to traverse the beam. Timing began once the mouse crossed onto the second (2.5cm) section of the beam and stopped once one of the forelimbs was inside the home cage. The maximum time of 60 seconds was recorded for animals that did not finish the assay within the allotted time or fell off the beam on the last (0.5cm) section. Test day trials were videotaped and later assessed in slow motion for number of steps and slips (errors) that occurred during the traversal. Slips were counted if at least ¾ of any limb left the beam. Errors per step were not recorded for animals that did not complete the task. For a detailed protocol, see <u>dx.doi.org/10.17504/protocols.io.kxygx4znzl8j/v1</u>.

For the pole descent assay, a 0.5m rod, 1cm in diameter, attached to a base and wrapped in shelf liner to increase grip, was used to assess the animal’s ability to climb down a pole. Mice were given 1 day of training prior to the test day, on the same days as beam training and testing, with about an hour of rest time between assays. The training day consisted of 3 trials. On the first trial, cagemates were removed and the pole was placed into the home cage. The animal was placed at the bottom of the home cage with the pole for approximately 1 minute to allow exploration. Then the animal was placed approximately ⅓ of the way up the pole, facing head down, and allowed to descend. For the second trial, the animal was placed ⅔ of the way up the pole, and for the third trial, at the top of the pole. On the consecutive day, animals were tested for their ability to descend from the top of the pole. Timing started at the moment the animal’s tail was released and ended when one of the hindlimbs touched the base of the pole. The maximum time of 60 seconds was recorded for animals that slid or fell off the pole apparatus. For a detailed protocol, see <u>dx.doi.org/10.17504/protocols.io.8epv5k9mjv1b/v1</u>.

For the wire hang assay, a clean cage with bedding was placed 40cm underneath a 30cm x 30cm wire mesh screen. Animals were placed at the center of the screen and gently flipped head-first until upside down. Timing started once the animal was hanging from the wire screen horizontally and stopped when the animal fell to the bedding underneath or remained hanging for 60 seconds. For detailed protocol, see <u>dx.doi.org/10.17504/protocols.io.6qpvrw62plmk/v1</u>.

For the adhesive removal assay, cagemates were removed from the home cage. A circular ⅜ inch sticker (Tough-Spots - Diversified Biotech/USA Scientific) was placed on the nose bridge of the animal being tested using tweezers. The mouse was released into the home cage and timing was started. Timing was stopped when the mouse successfully removed the sticker from its nose, with a maximum time of 30 seconds. For a detailed protocol, see <u>dx.doi.org/10.17504/protocols.io.4r3l21objg1y/v1</u>.

For the hindlimb clasping assay, mice were picked up by the tail, with the underside facing the observer, for approximately 10 seconds. Rigidity of the movement of hindlimbs was assessed on a scale of 0 to 3. A score of 0 was given if the animal was able to extend its hindlimbs outward freely, with flexibility and no clasping. A score of 1 was given if the animal had some tendency to clasp its hindlimbs inward, either one or both, but for the majority of the restraint the limbs were still spread outward and flexible. A score of 2 was given if hindlimbs were clasped inward for the majority of the time, but some movement and flexibility was still present. A score of 3 was given if the hindlimbs were clasped together entirely, with no movement or flexibility^45^. For a detailed protocol, see <u>dx.doi.org/10.17504/protocols.io.eq2ly4n2mlx9/v1</u>.

To measure fecal output, mice were individually placed into empty, bedding-free, standard mouse cages. The number of fecal pellets produced was counted after 15 minutes.

Following the fecal output assay, remaining fecal pellets were assessed according to the Bristol stool scale: a score of 1 was given for fecal pellets that were small, dry, hard lumps, indicating severe constipation. A score of 2 was given for lumpy, dry, but sausage-like fecal pellets, indicating mild constipation. A score of 3 was given for mildly dry sausage-like pellets with cracks in the surface. A score of 4 was given for smooth, soft, sausage or snake-shaped pellets. Both scores of 3 and 4 are considered normal. A score of 5 was given for soft blobs with clear-cut edges. A score of 6 was given for fluffy pieces with ragged edges, indicating mild diarrhea. Lastly, a score of 7 was given for an entirely liquid consistency, indicating severe diarrhea^135^. An average score was taken when animals produced fecal pellets of various scores during one fecal output session.

Following fecal output and score assays, 2-3 fecal pellets were collected in pre-weighed tubes. Tube and wet fecal pellet weights were measured and then pellets were incubated at 65℃ for 24 hours. Tubes and dry fecal pellets were re-weighed and percentage of water content calculated. For detailed protocols of constipation assays, see <u>dx.doi.org/10.17504/protocols.io.6qpvrw6qplmk/v1</u>.

To measure whole gut transit time, animals were gavaged with 150μl of autoclaved 6% (w/v) Carmine red (Millipore Sigma) and 0.5% methylcellulose (Sigma-Aldrich) solution in water. After 3 hours, animals were transferred to empty, bedding-free, standard mouse cages and observed every 30 minutes up to 8 hours for the first red fecal pellet^136,137^. For a detailed protocol, see <u>dx.doi.org/10.17504/protocols.io.rm7vz9yq8gx1/v1</u>.

For the bead expulsion assay, animals were placed into an isoflurane induction chamber on a heating pad in 1 L/min oxygen and a concentration of 3-5% isoflurane until the respiratory rate reached one breath per minute. Animals were then transferred to a nose cone to maintain anesthesia and a 2mm glass bead inserted into the colon 2 cm proximal to the anus using a metal gavage tube with lubricating jelly. Once the animal emerged from anesthesia, it was transferred to an individual empty, bedding-free, standard mouse cage and timed until the bead was expelled, up to 60 minutes^137,138^. For a detailed protocol, see <u>dx.doi.org/10.17504/protocols.io.n92ld6zn7g5b/v1</u>.

Motor and gastrointestinal function datasets in Figures 1 and 2 and Supplementary Figures 1 and 2 were analyzed using the Kruskal-Wallis test followed by a Conover-Iman post-hoc test of multiple comparisons with Benjamini-Hochberg false discovery rate (FDR)^139^ to correct for multiple testing. Graphs were generated using GraphPad Prism 10 software (http://www.graphpad.com, RRID:SCR_002798). P-values, n values, and standard errors are indicated in figures and legends.

For motor function datasets in Supplementary Figure 3, analysis was done using statistical models with either a simple right-censored (at 30 or 60 seconds depending on the test) LogNormal distribution or a mixture model that accounts for mice not engaging with the task within the allotted time. Analysis was performed using custom Python code (see Key Resources Table for GitHub repository of behavior analyses). All benCom-PD motor function assay data, as well as *F. prausnitzii* pole descent and adhesive removal assay results were fit into a simple LogNormal model:

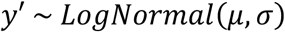

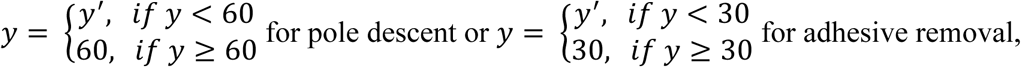

where 𝜇 (𝑚𝑢), 𝜎(𝑠𝑖𝑔𝑚𝑎) are the standard parameters of the LogNormal distribution.

*F. prausnitzii* beam traversal and wire hang assay data were fit into a right-censored LogNormal mixture model:\

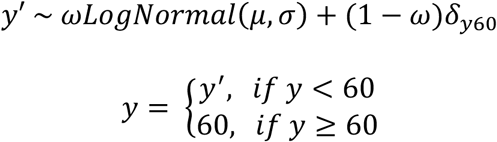

where 𝜔 (𝑜𝑚𝑒𝑔𝑎) is a parameter describing the probability of a subject mouse engaging with the test, 𝜇 (𝑚𝑢) and 𝜎(𝑠𝑖𝑔𝑚𝑎) are standard parameters of the LogNormal distribution, and 𝛿_60_ is the Kronecker delta function.

The models’ fits were evaluated with a graphical model assessment approach, using Q–Q plots and predictive ECDFs. Parameters were estimated by maximum likelihood, and 10,000 bootstrap replicates were generated to construct confidence intervals for group comparisons.

For datasets in Supplementary Figure 4, heterogeneity between the two *F*.

*prausnitzii* experimental cohorts was assessed using Cliff’s delta^140^ and its 95% confidence interval. Confidence intervals were estimated using bias-corrected and accelerated bootstrap resampling (50,000 iterations)^141^. For each bootstrap sample, Cliff’s delta was recalculated, and the 2.5th and 97.5th percentiles of the bootstrap distribution were taken as the confidence limits. Cochran’s Q^142^ and the I² statistic^143^ were assessed using the following formulas, assuming 1 degree of freedom:

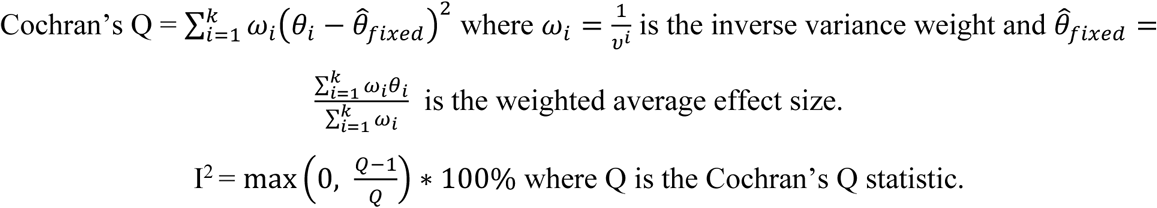

An I^2^ value ≤ 20% was taken as indicative of small to moderate heterogeneity across the two cohorts, based on all three pairwise group comparisons.

### Protein assays

Following motor and GI function testing, animals were sacrificed and tissues collected. For Western blot assays, 2mm sections of the substantia nigra and striatum were dissected, flash frozen, and stored at –80°C until further processing. Brain tissues were lysed using a handheld homogenizer (BT LabSystems) and the Bio-Plex Cell Lysis Kit according to manufacturer’s instructions (Bio-Rad). Protein concentrations were normalized using the Pierce BCA Protein Assay Kit (Thermo Scientific). Samples were separated on a 4-20% Tris-Glycine gel (Invitrogen) and transferred onto a 0.45µm PVDF membrane (Immobilon-P - Merck Millipore Ltd.). To enhance detection of αSyn, membranes were placed in 4% paraformaldehyde (PFA) solution for 30 minutes^144^. Blocking was performed using 5% bovine serum albumin (BSA) in Tris-buffered saline with 0.1% Tween-20 (TBS-T) for phosphorylated αSyn antibody or 5% nonfat milk in TBS-T for β-actin antibody. Membranes were cut at 25kDa and incubated overnight at 4℃ with primary antibodies: Anti-phosphorylated S129 α-Synuclein Rabbit (Abcam Cat# ab51253, RRID:AB_869973, 1:1000) or Anti-β-actin Mouse (Abcam Cat# ab8226, RRID:AB_306371, 1:1000) in the respective blocking buffers. After washing, membranes were incubated with Anti-IgG HRP rabbit or mouse antibodies (Cell Signaling Technology Cat# 7074, RRID:AB_2099233 and Cat# 7076, RRID:AB_330924, 1:1000) in TBS-T for 1.5 hours at room temperature. Signal was detected using the Clarity Western ECL Substrate (Bio-Rad) on a Bio-Rad ChemiDoc imaging system. Densitometry analysis was performed using ImageJ. For a detailed protocol, see <u>dx.doi.org/10.17504/protocols.io.x54v92z2ql3e/v1</u>.

For dot blot, homogenized brain tissue samples were diluted to 500ng/μL of protein and directly spotted onto a 0.45um nitrocellulose membrane (Thermo Scientific). Membranes were blocked using 5% nonfat milk in TBS-T and incubated for 1.5 hours with Anti-Aggregated α−Synuclein Rabbit (Abcam Cat# ab209538, RRID:AB_2714215, 1:1000) at room temperature in blocking buffer. Subsequent steps followed the same procedure as described for Western blot analysis. For a detailed protocol, see <u>dx.doi.org/10.17504/protocols.io.261gen2xdg47/v1</u>.

For multiplex bead assay, proximal large intestine segments (∼2 cm) were homogenized in lysis buffer (Bio-Plex Cell Lysis Kit, Bio-Rad) using Lysing Matrix D tubes and a bead beater (MP Biomedicals). Total protein concentrations were quantified and normalized using the Pierce BCA Protein Assay Kit (Thermo Scientific). Cytokine levels were then measured on the BioPlex 200 platform using the Bio-Plex Pro Mouse Th17 6-plex Assay (Bio-Rad), following the manufacturer’s protocol.

### Flow cytometry

Spleen and mesenteric lymph nodes were collected and processed for flow cytometric analysis. Tissues were passed through 70µm cell strainers, and spleen cells were treated with red blood cell lysing buffer (Hybri-Max - Sigma Aldrich) for 5 minutes at room temperature. For a detailed isolation protocol, see <u>dx.doi.org/10.17504/protocols.io.nm2dc8e.</u> For later intracellular staining, isolated cells were stimulated with phorbol myristate acetate (PMA) (50 ng/ml, Sigma Aldrich) and ionomycin (750 ng/ml, Sigma Aldrich) with GolgiPlug^TM^ (1:1000, BD Biosciences) for 4 hours at 37°C and 5% CO_2_. All cells were then blocked using anti-Fcy receptor (anti-CD16/CD32) (clone 93, eBioscience) and labelled with the following fluorescent-conjugated antibodies: CD45 (clone 30-F11, eBioscience), TCRB (clone H57-597, eBioscience), CD4 (clone RM4-5, eBioscience), and CD25 (clone PC61.5, eBioscience) (all at 1:500 concentration). All cells were labelled with Fixable Near-IR LIVE/DEAD Stain Kit (Invitrogen) to distinguish live cells from dead/debris. For intracellular staining, cells were treated using the Foxp3/Transcription Factor Fixation/Permeabilization Kit (eBioscience) according to the manufacturer’s instructions and stained with fluorescent-conjugated antibody against FoxP3 (clone FJK-16s, eBioscience, RRID:AB_763537) at 1:100 concentration. Data were acquired on a CytoFLEX S Analyzer (Beckman Coulter) and analyzed with FlowJo software (BD, Tree Star, https://www.flowjo.com/solutions/flowjo, RRID:SCR_008520).

Data generated from protein and cellular assays was analyzed using a one-way ANOVA with Tukey post-hoc test and graphs generated with GraphPad Prism 10 software. P–values, n values, and standard errors are indicated in figures and legends.

### RNA sequencing & qPCR analysis

Approximately 1cm of proximal large intestine was collected, immediately flash-frozen in TRIzol reagent (Zymo Research), and stored at –80°C. Total RNA was extracted using the Direct-zol RNA MiniPrep Plus Kit (Zymo Research) per manufacturer’s protocol, including an on-column DNase I treatment. RNA quality and concentration were assessed using a NanoDrop spectrophotometer and an Agilent 2100 Bioanalyzer (Agilent Technologies). Libraries were prepared using the ABclonal FAST RNA-seq Library Prep Kit. Pooled libraries underwent 50 base pair, paired-end sequencing at a target depth of 30 million reads per sample on the Illumina NovaSeq X platform.

Raw sequencing reads were aligned to the *Mus musculus* GRCm36 Primary Assembly reference genome (Release M36 from GENCODE) using STAR aligner v2.7.1 (https://github.com/alexdobin/STAR, RRID:SCR_004463)^145^ with default parameters. Gene-level read counts were quantified using featureCounts^146^. Only protein coding genes with a minimum sum of at least 10 counts across all samples were retained. Normalization and differential gene expression analysis was performed using the DESeq2 package (https://bioconductor.org/packages/release/bioc/html/DESeq2.html, RRID:SCR_015687)^147^ in R (v4.3.0, http://www.r-project.org, RRID:SCR_001905). Genes with an adjusted p-value < 0.05 and log2 fold change were considered significantly differentially expressed. Gene ontology and pathway enrichment analyses for differentially expressed genes were conducted using the clusterProfiler package (http://bioconductor.org/packages/release/bioc/html/clusterProfiler.html, RRID:SCR_016884)^148^. Gene sets with an adjusted p-value < 0.1 and |log_2_ fold-change| > 0.2 were analyzed for up and down regulated genes separately for each comparison of interest. Data visualization, including volcano plots and principal component analysis (PCA), was performed using ggplot2 (https://cran.r-project.org/web/packages/ggplot2/index.html, RRID:SCR_014601)^149^.

### Volatile fatty acid measurement

Fecal pellets, cecal contents, and serum were collected and flash frozen on dry ice for later processing. Fecal pellets were collected fresh from live animals at 18 weeks of age. Cecal contents were collected at 23 weeks during tissue collection. For serum isolation, blood was drawn via cardiac puncture or from the trunk into microtubes containing serum gel (Z-gel, Sarstedt AG & Co.), allowed to clot on ice for 30 minutes, then centrifuged at 10,000xg for 5 minutes. The resulting serum was collected for downstream analysis.

Fecal, cecal, and blood serum volatile fatty acid (VFA) concentrations were quantified by high-performance liquid chromatography (HPLC) using sample extracts. For fecal and cecal extracts, deionized water was added to the samples at a ratio of 1 mL of water to 100 mg of sample. For serum extracts, in order to precipitate any proteins in the serum, 2.5 mM H_2_SO_4_ was added to samples at a 1:1 ratio. Serum samples were then vortexed briefly and placed at 4°C for 20 minutes. All extracts were vortexed at 3,200 rpm for 5 minutes and then centrifuged at 13,000 rpm for 15 minutes at 4°C. The supernatants were filter-sterilized through a 0.22 μm syringe filter. The filtrates were used for HPLC analysis and soluble chemical oxygen demand (sCOD) measurements. Analysis of VFAs was performed on an Agilent 1260 Infinity II (Agilent, Santa Clara, California, USA) diode array detector at 210 nm with a Bio-Rad Aminex HPX-87H column (Cat. #: 1250095) as previously described^150,151^. Samples were analyzed at a column temperature of 65°C using 5 mM H_2_SO_4_ as a mobile phase at a flow rate of 0.6 mL/min for 80 minutes. A standard curve was generated using a 10 mM VFA mixed standard (Supelco Volatile Free Acid Mix, Cat. #: CRM46975) containing acetate, formate, butyrate, isobutyrate, valerate, isovalerate, propionate, caproate, and heptanoate. Dilutions were done in triplicate to achieve the following concentrations: 0.25 mM, 2 mM, 5 mM, 7.5 mM, and 10 mM. To account for differences in sample consistency, all VFA measurements were normalized against the sCOD measured from the filtrate, as previously reported^95,152^. To measure sCOD, a HACH high-range (20 – 1500 mg COD/L) COD kit (Cat. #: 2565115) was used^152–154^. 100 μL of fecal or cecal sample filtrate, or 25 μL of serum sample filtrate was transferred to a HACH tube and adjusted to a final volume of 2 mL with deionized water. Samples were digested on a HACH DRB200 Digital Reactor Block (Cat. #: LTV082.53.44001) according to the manufacturer’s recommendations. A HACH DR6000 spectrophotometer (Cat. #: LPV441.99.00012) was used to measure sCOD. Resultant data was analyzed using a one-way ANOVA with Tukey post-hoc test and graphs generated with GraphPad Prism 10 software. P-values, n values, and standard errors are indicated in figures and legends.

### Fecal microbiome analysis with shotgun metagenomics

Fecal pellets for metagenomics were collected fresh from live animals at 18 weeks of age and DNA was isolated from one mouse stool sample pellet using the Qiagen QIAamp PowerFecal Pro kit following the manufacturer’s instructions. Aliquots were shipped to Prebiomics S.r.l. (Trento, Italy) for library preparation and sequencing.

DNA was quantified using the Quant-iT™ 1X dsDNA Assay Kits, BR (Life Technologies) in combination with the Varioskan LUX Microplate Reader (Thermo Fisher Scientific). Sequencing libraries were prepared with the Illumina DNA Prep (M) Tagmentation (96 Samples, IPB) kit in combination with Illumina DNA/RNA UD Indexes. Amplified libraries were purified using the double-sided bead purification procedure, as described in the Illumina protocol. Then, library concentrations (ng/µL) were quantified and base pair lengths (bp) evaluated using the D5000 ScreenTape Assay in combination with TapeStation 4150 (Agilent Technologies) to establish library pooling volumes. Pooling was then quantified with the Qubit 1x dsDNA HS kit (Life Technologies) and a Qubit 3.0 Fluorometer (Life Technologies). Libraries were sequenced using the Novaseq X Plus platform (Illumina) at an average depth of 15 billion base pairs per sample. For a detailed protocol, see <u>dx.doi.org/10.17504/protocols.io.e6nvw15dwlmk/v1</u>.

FASTQ files were uniformly processed using the curated curatMetagenomicsNextflow pipeline (https://github.com/seandavi/curatedMetagenomicsNextflow) (commit 4cde6fb). Mouse genomic reads were removed in the KneadData step using the C57BL mouse reference genome. The core output of the pipeline includes quality control (FastQC and Trimmomatic via KneadData) and MetaPhlAn v4.1.1 (https://huttenhower.sph.harvard.edu/metaphlan/, RRID:SCR_004915)^155^ and HUMANnN v3.9 (https://huttenhower.sph.harvard.edu/humann, RRID:SCR_014620)^156^ abundance profiles, which were used for statistical analyses. All analyses were performed in R 4.5.1. System, R, and package versions were reported with *sessionInfo()* at the end of all analysis scripts (see Data and Materials Availability).

Data were downloaded and formatted for analyses using the R package parkinsonsMetagenomicData (https://github.com/ASAP-MAC/parkinsonsMetagenomicData). Samples were analyzed at the species genome bin (SGB) level except for relative abundance stackplots (Fig. 3A), which were aggregated at the phylum level to aid legend readability.

The mia package (https://bioconductor.org/packages/release/bioc/html/mia.html, RRID:SCR_023619) was used to perform alpha and beta diversity analyses. Observed richness and Shannon diversity were calculated with addAlpha. ANOVA was used to test for differences in means between genotype-treatment groups. The Bray-Curtis dissimilarity matrix was calculated with addDissimilarity on relative abundance data ranging from 0 to 1. The function addMDS coloring by genotype-treatment was used to plot the first two axes of the principal coordinate analysis (PCoA) on the Bray-Curtis dissimilarity matrix. To test for genotype-treatment effect on the overall microbiome composition, a permutational multivariate analysis of variance (PERMANOVA)^157^ followed by permutational test for group homogeneity (permutest) was used, both implemented in addPERMANOVA. To further test for non-overlap between groups, a pairwise PERMANOVA with pairwiseAdonis was used, a custom function documented at https://github.com/pmartinezarbizu/pairwiseAdonis. False Discovery Rate (FDR) was calculated for all pairwise tests on beta-diversity (e.g., 3 pairwise tests for 3 groups) using the Benjamini-Hochberg method.

The effect of probiotic treatment on the relative abundance and prevalence of each SGB was determined using MaAsLin3^158^. SGB relative abundances in each sample were scaled from 0 to 1, followed by a log2 transformation. Thy-1-ASO treated mice were compared to Thy-1-ASO control mice. False discovery rate across all SGBs was calculated using the Benjamini-Hochberg method as implemented in MaAsLin3.

For mediation analysis, behavior data from only one *F. prausnitzii* cohort was used to match sequencing data. The first three principal coordinates were used as continuous mediators using the Baron and Kenny approach^159^, with the mediate function implemented in the “mediation” R package (https://cran.r-project.org/web/packages/mediation/index.html, RRID:SCR_026984). Confidence intervals were estimated using quasi-Bayesian simulations (n = 1000)^160^.

### Data and Materials Availability

The data, code, protocols, and key lab materials used and generated in this study are listed in the Key Resource Table (**Supplementary Table 2**) alongside their persistent identifiers. For microbiome analysis data, raw FASTQ files were deposited in the SRA (PRJNA1259538) along with sample metadata, and code to generate data, figures, and statistics is available at https://doi.org/10.5281/zenodo.17128213. For RNA sequencing analysis data, raw FASTQ files were deposited in the SRA (PRJNA1308739) along with sample metadata, and code to generate data, figures, and statistics is available at https://github.com/jboktor/fprausnitzii-treatment-rnaseq. Raw flow cytometry data were deposited to Zenodo (https://doi.org/10.5281/zenodo.16929959) along with sample metadata. All other raw data and analyses, including behavior data analysis, can be found at https://doi.org/10.5281/zenodo.17137678.

### Author Contributions

A.M. and S.K.M. designed the studies and wrote the manuscript. A.M. led all investigations and drafted figures. Metagenomic analysis and figure creation was executed by G.A., while data processing was done by K.L., with supervision from N.S. and L.D.W. A.M.S. assisted with flow cytometry experiments, tissue collections, and initial manuscript editing. J.C.B. carried out downstream RNA sequencing analysis and figure creation for this data. B.D. collected volatile fatty acid measurements with assistance from D.D., and supervision from R.K.B. A.D.O. performed statistical analysis for motor and GI function assays. A.V.W. supported the design and quality control of RNA sequencing methodology, and raw data processing, with supervision from R.F.I. P.S. assisted with behavior data collection and tissue collections. All authors read and approved the final manuscript.

## Acknowledgements

The authors would like to thank the members of the Mazmanian laboratory for their helpful critiques and review of the manuscript. We thank T. Thron for husbandry and maintenance of the Thy1-ASO “Line 61” mouse line, and L.B. De los Santos, I. Cardenas, and J. Gutierrez for animal care. We are grateful to C. Oikonomou for manuscript editing and submission support, and Y. Garcia-Flores for lab administrative support. We would also like to thank J. Griffiths for help with tissue collections. Flow cytometric analysis was performed at the Caltech Flow Cytometry and Cell Sorting Facility, with support from M. Gregory. Multiplex analysis was done at the Caltech Protein Expression Center, with support from M. Anaya. RNA sequencing was carried out by the UCLA Technology Center for Genomics & Bioinformatics (TCGB).

Metagenomics sequencing was made possible by Prebiomics S.r.l. (Trento, Italy), with support from M. Bolzan. Figures 1A, B and 2A were created in BioRender (Moiseyenko, A., 2025, https://BioRender.com/zvkyy6g). This research was funded in part by Aligning Science Across Parkinson’s (ASAP-020495 and ASAP-000375) through the Michael J. Fox Foundation for Parkinson’s Research (MJFF), as well as the Heritage Medical Research Institute to S.K.M. For the purpose of open access, the authors have applied a CC BY public copyright license to all Author Accepted Manuscripts arising from this submission.

