## Supplementary Information for "*Faecalibacterium prausnitzii*, depleted in the Parkinson’s disease microbiome, improves motor deficits in α-synuclein overexpressing mice"

for

Anastasiya Moiseyenko *et al.*

**This file includes:**

- Supplementary Figures 1 through 8
- Supplementary Tables 1 and 2

**Other supplementary materials provided in separate files:**

- Supplementary Data

#### Supplementary Figure 1

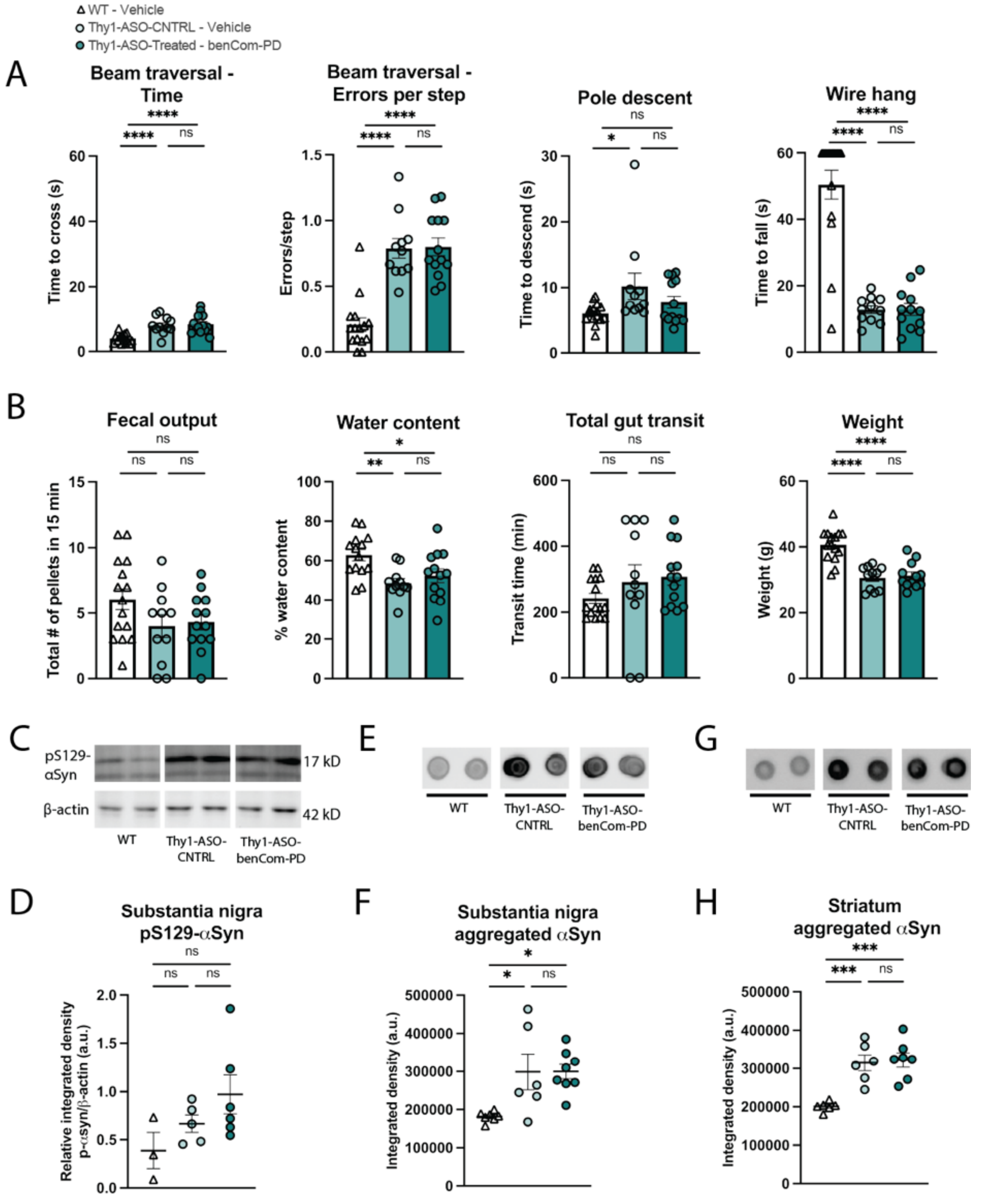

**Supplementary Figure 1**. **Motor and GI function tests and α-synuclein assays not altered by benCom-PD treatment.** *(Associated with Figure 1)*. (**A**) Motor function and (**B**) GI function and animal weights of benCom-PD-treated Thy1-ASO animals and controls tested at 20 weeks. N = 11-15. Points represent individual animals; bars represent standard error of the mean. (**C**)

Representative images of Western blots of phosphorylated-S129-α-synuclein in substantia nigra and (**D**) quantification. N = 3-6. (**E-H**) Representative images of dot blots for aggregated α-synuclein in (E) substantia nigra and (G) striatum and (F, H) quantification. N = 6-8. Points represent individual animals; bars represent standard error of the mean. Behavior data in (A, B) analyzed by Kruskal-Wallis test followed by the Conover-Iman post-hoc test with Benjamini-Hochberg false discovery rate (FDR) correction; protein data in (D-H) analyzed by one-way ANOVA with Tukey post-hoc test. ns – not significant; *p≤0.05; **p≤0.01; ***p≤0.001; ****p ≤0.0001.

Abbreviations: WT – wildtype; Thy1-ASO-CNTRL – Thy1-human α-synuclein overexpressing; αSyn – α-synuclein; benCom – beneficial commensal consortium.

**Supplementary Figure 2**

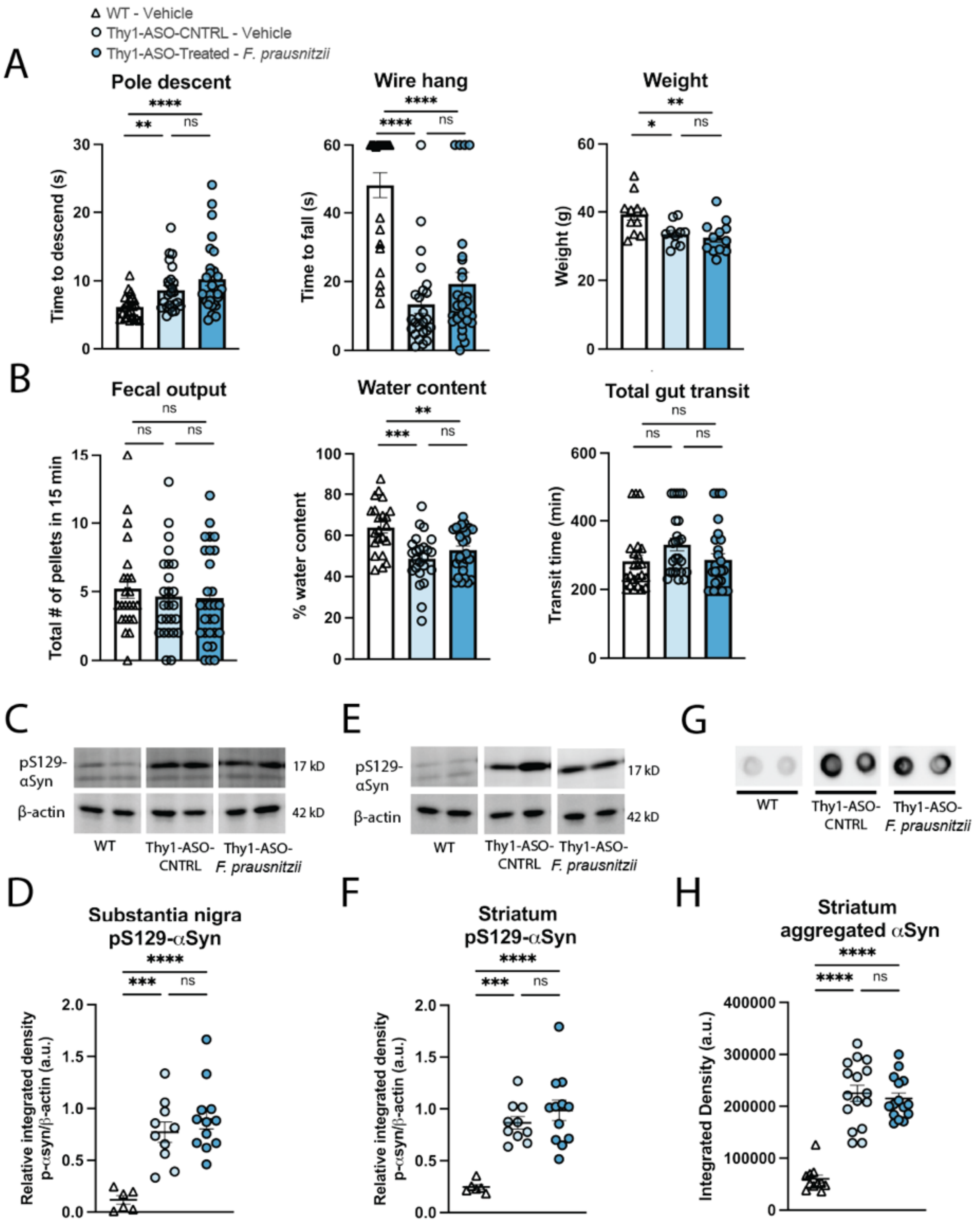

**Supplementary Figure 2. Motor and GI function tests and αSyn assays not altered by *F. prausnitzii* treatment.** *(Associated with Figure 2)* (**A**) Motor function and animal weights, and (**B**) GI function for *F. prausnitzii*-treated Thy1-ASO animals and controls tested at 20 weeks. N = 23-29. Points represent individual animals, compiled from two independent cohorts; bars represent standard error of the mean. (**C-F**) Representative images (C, E) and quantification (D, F) of Western blots for phosphorylated-S129-α-synuclein in (C, D) substantia nigra and (E, F)

striatum. N = 6-12. (**G, H**). Representative images (G) of dot blots and quantification (H) for aggregated α-synuclein in the striatum. N = 11-16. Points represent individual animals, compiled from two independent cohorts; bars represent standard error of the mean. Behavior data in (A, B) analyzed by Kruskal-Wallis test followed by the Conover-Iman post-hoc test with Benjamini-Hochberg false discovery rate (FDR) correction; protein data in (C-H) analyzed by one-way ANOVA with Tukey post-hoc test. ns – not significant; *p≤0.05; **p≤0.01; ***p≤0.001; ****p ≤0.0001.

Abbreviations: WT - wildtype, Thy1-ASO-CNTRL – Thy1-human α-synuclein overexpressing control; pS129 αSyn – phosphorylated-S129-α-synuclein; αSyn – α-synuclein

#### Supplementary Figure 3

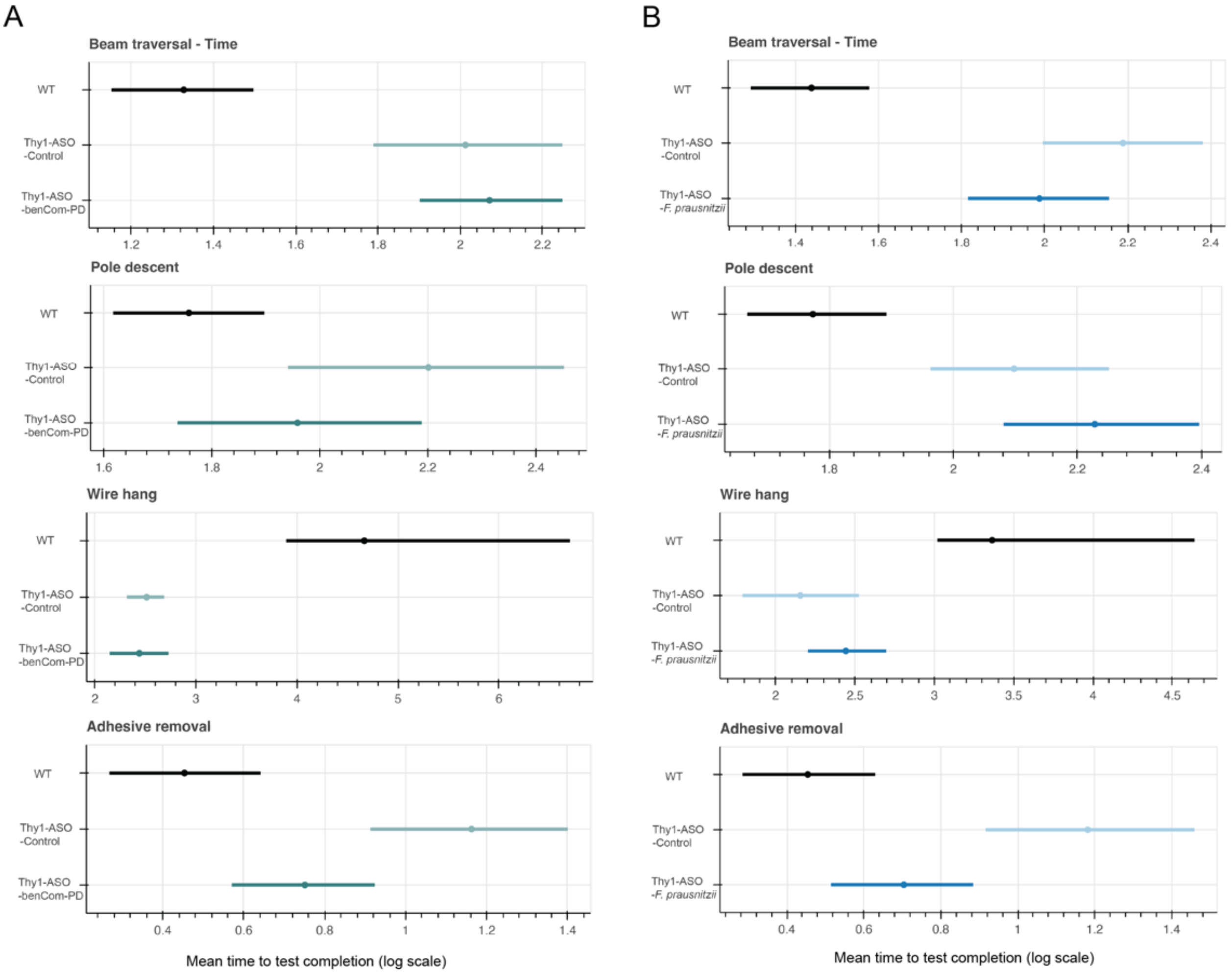

**Supplementary Figure 3. Parametric inference analysis of motor function tasks.** *(Associated with Figure 2)* Performance of mice treated with (**A**) benCom-PD and (**B**) *F. prausnitzii* was analyzed using generative statistical models that account for ceilings in task completion. Data consisted of individual animals from a single cohort (A) or two independent cohorts (B). Confidence intervals (95% CI) were calculated using maximum likelihood estimation (MLE) followed by parametric bootstrap. The μ 95% CI parameter describes the mean test completion time displayed on a logarithmic scale.

Abbreviations: WT - wildtype, Thy1-ASO – Thy1-human α-synuclein overexpressing

#### Supplementary Figure 4

A

| Test | Group | Q | I² (%) |
|---|---|---|---|
| Beam traversal - Time | Thy1-ASO-Cntrl vs.-Fp | 0.426508 | 0 |
| | WT vs. Thy1-ASO-Fp | 1.067422 | 6.316368 |
| | WT vs. Thy1-ASO-Cntrl | 0.121443 | 0 |
| Beam traversal – Errors per step | Thy1-ASO-Cntrl vs.-Fp | 1.961784 | 49.02599 |
| | WT vs. Thy1-ASO-Fp | 0.004704 | 0 |
| | WT vs. Thy1-ASO-Cntrl | undefined | 0 |
| Pole descent | Thy1-ASO-Cntrl vs.-Fp | 1.038982 | 3.751952 |
| | WT vs. Thy1-ASO-Fp | 0.068595 | 0 |
| | WT vs. Thy1-ASO-Cntrl | 0.65088 | 0 |
| Wire hang | Thy1-ASO-Cntrl vs.-Fp | 0.00241 | 0 |
| | WT vs. Thy1-ASO-Fp | 0.249885 | 0 |
| | WT vs. Thy1-ASO-Cntrl | 0.997635 | 0 |
| Adhesive removal | Thy1-ASO-Cntrl vs.-Fp | 0.008109 | 0 |
| | WT vs. Thy1-ASO-Fp | 0.008606 | 0 |
| | WT vs. Thy1-ASO-Cntrl | 1.11974 | 0 |
| Hindlimb clasp | Thy1-ASO-Cntrl vs.-Fp | 0.458192 | 0 |
| | WT vs. Thy1-ASO-Fp | 3.418827 | 70.750201 |
| | WT vs. Thy1-ASO-Cntrl | 0.614324 | 0 |

B

| Test | Group | Q | I² (%) |
|---|---|---|---|
| Fecal output | Thy1-ASO-Cntrl vs.-Fp | 1.698515 | 41.125055 |
| | WT vs. Thy1-ASO-Fp | 0.179903 | 0 |
| | WT vs. Thy1-ASO-Cntrl | 1.475329 | 32.218513 |
| Fecal score | Thy1-ASO-Cntrl vs.-Fp | 4.719061 | 78.809343 |
| | WT vs. Thy1-ASO-Fp | 1.506637 | 33.627016 |
| | WT vs. Thy1-ASO-Cntrl | 6.423796 | 84.432881 |
| Water content | Thy1-ASO-Cntrl vs.-Fp | 0.91584 | 0 |
| | WT vs. Thy1-ASO-Fp | 3.229106 | 69.031675 |
| | WT vs. Thy1-ASO-Cntrl | 3.791861 | 73.627726 |
| Total gut transit | Thy1-ASO-Cntrl vs.-Fp | 6.398381 | 84.371047 |
| | WT vs. Thy1-ASO-Fp | 5.014129 | 80.056356 |
| | WT vs. Thy1-ASO-Cntrl | 0.862981 | 0 |
| Bead expulsion | Thy1-ASO-Cntrl vs.-Fp | 1.503285 | 33.479015 |
| | WT vs. Thy1-ASO-Fp | 0.00756 | 0 |
| | WT vs. Thy1-ASO-Cntrl | 3.490956 | 71.354553 |

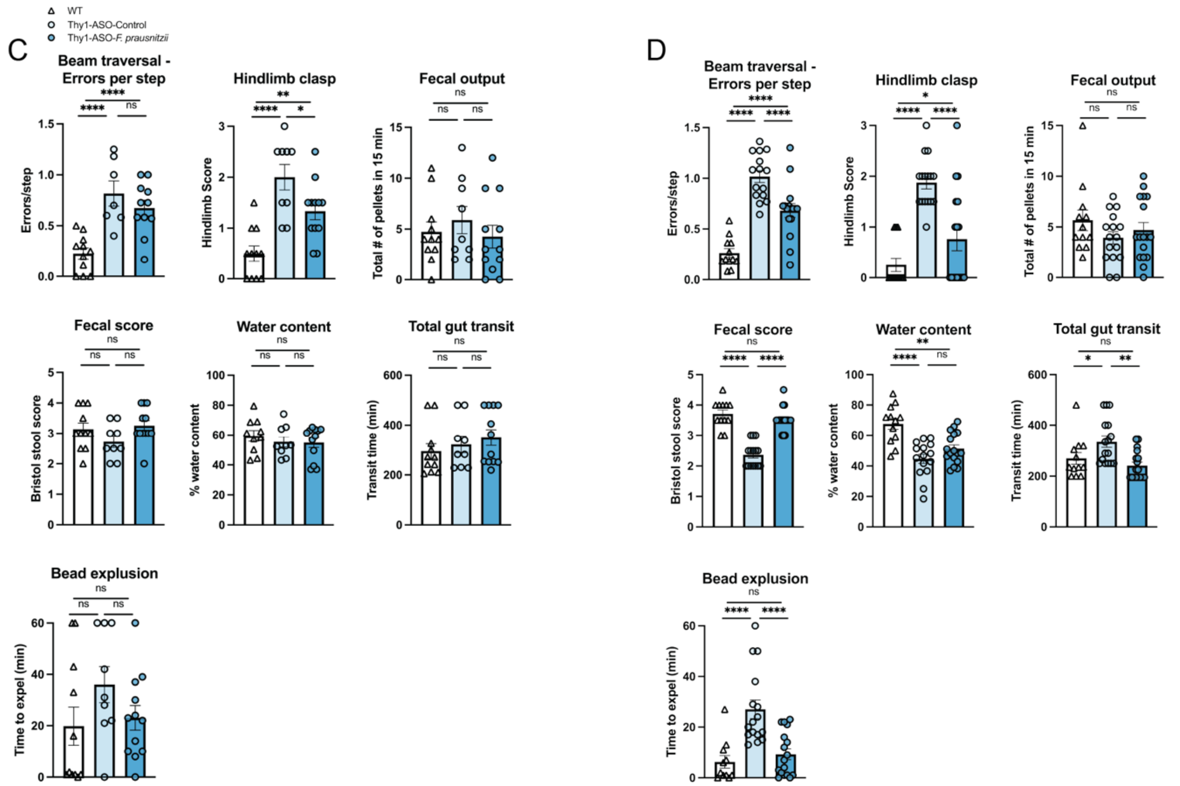

**Supplementary Figure 4. Analysis of heterogeneity across cohorts for motor and GI function assays in animals treated with *F. prausnitzii*.** *(Associated with Figure 2)* **(A, B)** Variability between cohorts, quantified by Cochran's Q and $I^2$ statistics, for pairwise comparisons of (A) motor and (B) gut physiology tests. An $I^2$ value ≤ 20% indicates low to moderate heterogeneity. (**C, D**) Individual cohort results for comparisons with higher heterogeneity: cohort 1, (C, N = 7-11) and cohort 2 (D, N = 12-17). Points represent individual animals; bars represent standard error of the mean. ns – not significant; *$p \leq 0.05$; **$p \leq 0.01$; ***$p \leq 0.001$; ****$p \leq 0.0001$.

Abbreviations: WT - wildtype, Thy1-ASO-CNTRL – Thy1-human α-synuclein overexpressing control; Fp – *F. prausnitzii*

#### Supplementary Figure 5

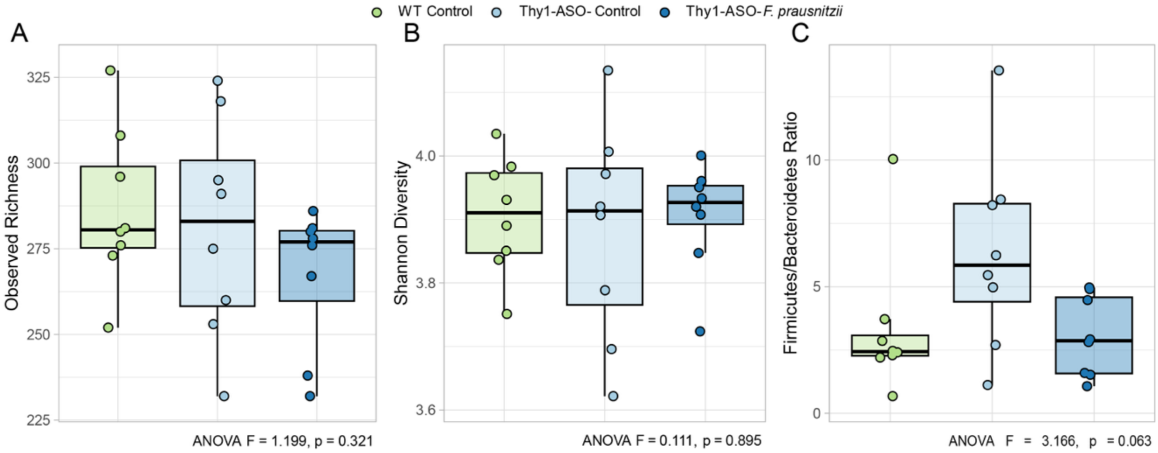

**Supplementary Figure 5. Gut microbiome aspects unaltered by *F. prausnitzii* treatment.** *(Associated with Figure 3)*. Distributions of (**A**) observed fecal SGB richness, (**B**) Shannon diversity, and (**C**) Firmicutes/Bacteroidetes ratio. Data were analyzed using a one-way ANOVA.

Abbreviations: WT- wildtype; Thy1-ASO – Thy1-human α-synuclein overexpressing; *F. prausnitzii – Faecalibacterium prausnitzii*

**Supplementary Figure 6**

A

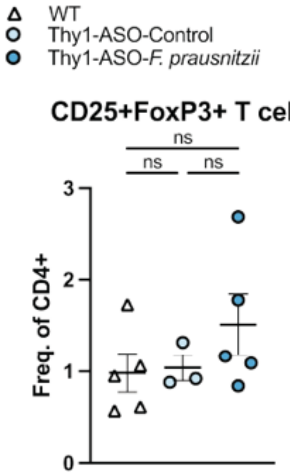

**Supplementary Figure 6. T$_{REGS}$ not altered by *F. prausnitzii* supplementation**. *(Associated with Figure 4)*. (**A**) Quantification of regulatory T cells in mesenteric lymph nodes, expressed as frequency of CD4+ cells. N = 3-5. Points represent individual animals, bars represent standard error of the mean. Data analyzed by one-way ANOVA with Tukey post-hoc test.

Abbreviations: WT – wildtype; Thy1-ASO-CNTRL – Thy1-human α-synuclein overexpressing control; *F. prausnitzii – Faecalibacterium prausnitzii*; ns – not significant

**Supplementary Figure 7**

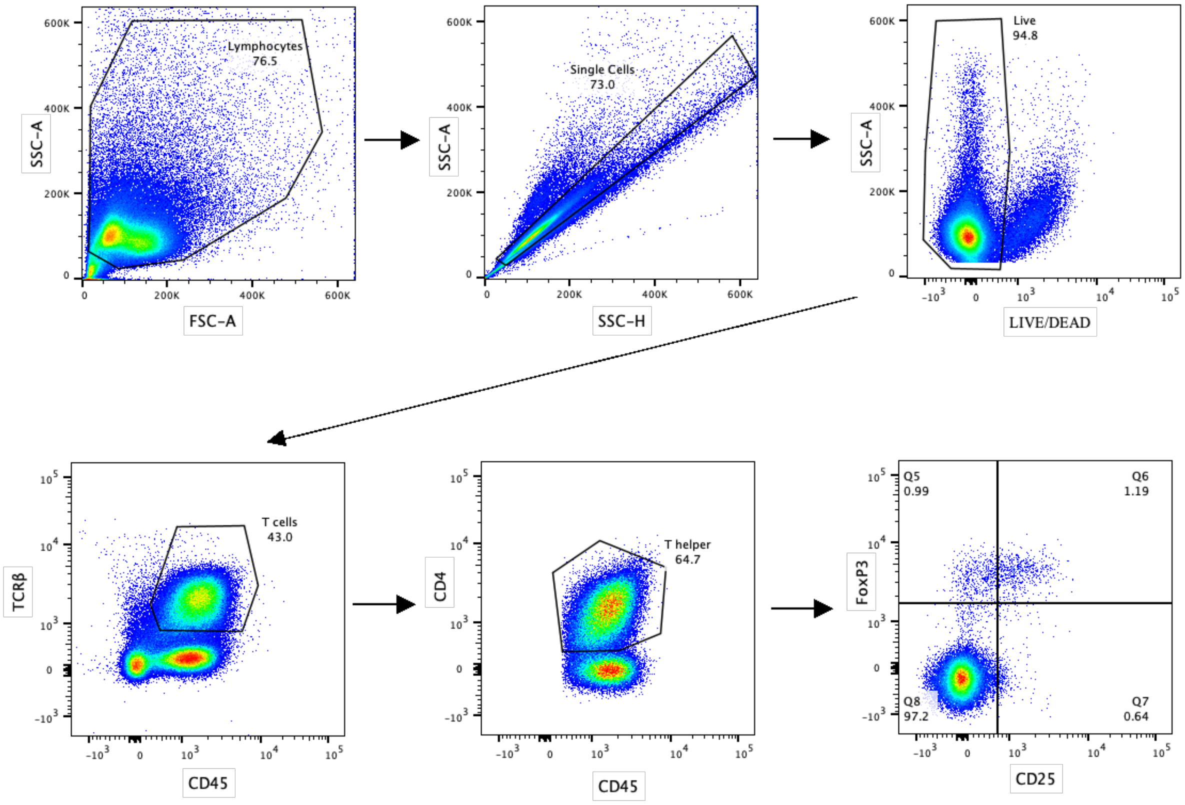

**Supplementary Figure 7. Adaptive immune system gating strategy for flow cytometry analysis.** *(Associated with Figure 4)*

### Supplementary Figure 8

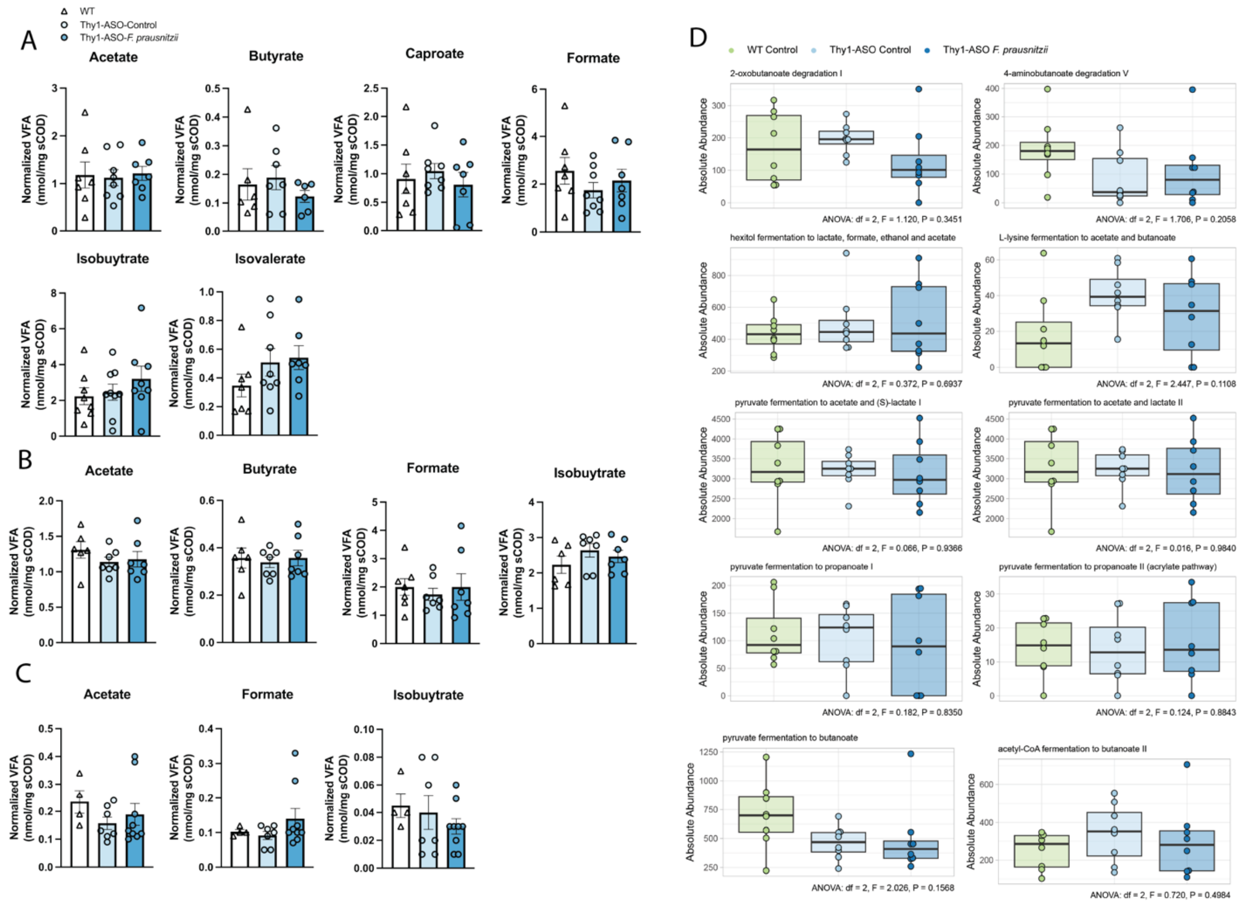

**Supplementary Figure 8. Volatile fatty acid and fermentation pathway analyses.** (**A**) Fecal, (**B**) cecal, and (**C**) serum detectable volatile fatty acid measurements, normalized by soluble chemical oxygen demand (sCOD), from Thy1-ASO animals treated with *F. prausnitzii* and controls. N = 4-9. Points represent individual animals, bars represent standard error of the mean. Data analyzed by one-way ANOVA with Tukey post-hoc test. No significant changes were detected. Points represent individual animals, bars represent standard error of the mean. (**D**) Boxplots of the abundance of predicted fermentation pathways inferred from metagenomic sequencing data using HUMAnN3. Boxes indicate the median and interquartile range. Data were analyzed using a one-way ANOVA.

Abbreviations: WT – wildtype; Thy1-ASO – Thy1-human α-synuclein overexpressing; *F. prausnitzii – Faecalibacterium prausnitzii*

#### Supplementary Table 1

| **Taxon Name** | **Taxonomic Level** | **Total signatures** | **Increased signatures** | **Decreased signatures** |
|---|---|---|---|---|
| *Akkermansia* | genus | 37 | 37 | 0 |
| *Faecalibacterium* | genus | 46 | 4 | 42 |
| *Roseburia* | genus | 45 | 4 | 41 |
| *Bifidobacterium* | genus | 33 | 30 | 3 |
| *Blautia* | genus | 25 | 2 | 23 |
| *Faecalibacterium prausnitzii* | species | 16 | 0 | 16 |
| *Fusicatenibacter* | genus | 19 | 1 | 18 |
| *Roseburia intestinalis* | species | 12 | 0 | 12 |
| *Peptoniphilus* | genus | 12 | 12 | 0 |
| *Alistipes* | genus | 12 | 12 | 0 |

**Supplementary Table 1**. **Summary of the top 10 microbial signatures across human Parkinson's disease studies.** Top 10 taxa found to be altered at the genus and species levels as of September 11, 2025, using BugSigDB (https://bugsigdb.org/; Geistlinger et al., 2024), a comprehensive database of published microbial signatures. Refer to **Supplementary Data 1** for an extended list of signatures.

**Supplementary Table 2. Key Resource Table**
Data, code, protocols, and key lab materials used and generated in this study.

| RESOURCE TYPE | RESOURCE NAME | SOURCE | IDENTIFIER | NEW/ REUSE | ADDITIONAL INFORMATION |
|---|---|---|---|---|---|
| Antibody | Anti-phosphorylated S129 a-Synuclein Rabbit | abcam | ab51253; RRID:AB_869973 | Reuse | 1:1000 concentration |
| Antibody | Anti-β-actin Mouse | abcam | ab8226; RRID:AB_306371 | Reuse | 1:1000 concentration |
| Antibody | Anti-IgG HRP Rabbit | Cell Signaling Technology | 7074; RRID:AB_2099233 | Reuse | 1:1000 concentration |
| Antibody | Anti-IgG HRP Mouse | Cell Signaling Technology | 7076; RRID:AB_330924 | Reuse | 1:1000 concentration |
| Antibody | Anti-Aggregated a-Synuclein Rabbit | abcam | ab209538; RRID:AB_2714215 | Reuse | 1:1000 concentration |
| Antibody | anti-Fcγ receptor (anti-CD16/CD32) (clone 93) | eBioscience | 14-0161-86; RRID:AB_467135 | Reuse | 1:100 concentration |
| Antibody | CD45 (clone 30-F11) | eBioscience | 48-0451-82; RRID:AB_1518806 | Reuse | 1:500 concentration |
| Antibody | TCRB (clone H57-597) | eBioscience | 45-5961-82; RRID:AB_925763 | Reuse | 1:500 concentration |
| Antibody | CD4 (clone RM4-5) | eBioscience | 25-0042-82; RRID:AB_469578 | Reuse | 1:500 concentration |
| Antibody | CD25 (clone PC61.5) | eBioscience | 17-0251-82; RRID:AB_469366 | Reuse | 1:500 concentration |
| Antibody | FoxP3 (clone FJK-16s) | eBioscience | 53-5773-82; RRID:AB_763537 | Reuse | 1:100 concentration |
| Bacterial strain | *Roseburia intestinalis* | DSMZ | DSMZ14610 | Reuse | |
| Bacterial strain | *Roseburia faecis* | DSMZ | DSMZ16840 | Reuse | |
| Bacterial strain | *Anaerostipes hadrus* | ATCC | ATCC29173 | Reuse | |
| Bacterial strain | *Fusicatenibacter saccharivorans* | DSMZ | DSMZ26063 | Reuse | |
| Bacterial strain | *Prevotella histicola* | DSMZ | DSMZ19854 | Reuse | |
| Bacterial strain | *Bacteroides ovatus* | ATCC | ATCC8483 | Reuse | |

| | | | | | |
|---|---|---|---|---|---|
| Bacterial strain | *Eubacterium rectale* | ATCC | ATCC33656 | Reuse | |
| Bacterial strain | *Faecalibacterium prausnitzii* | DSMZ | DSMZ17677 | Reuse | |
| Chemical, peptide, or recombinant protein | Carmine | Millipore Sigma | C1022 | Reuse | |
| Chemical, peptide, or recombinant protein | Phorbol Myristate Acetate (PMA) | Sigma Aldrich | P1585 | Reuse | 50 ng/ml |
| Chemical, peptide, or recombinant protein | Ionomycin | Sigma Aldrich | I0634 | Reuse | 750 ng/ml |
| Critical commercial assay | Pierce BCA Protein Assay Kit | Thermo Scientific | 23225 | Reuse | |
| Critical commercial assay | Bio-Plex Pro Mouse Th17 6-plex Assay | Bio-Rad | M6000007NY | Reuse | |
| Critical commercial assay | Direct-zol RNA MiniPrep Plus Kit | Zymo Research | R2072 | Reuse | |
| Critical commercial assay | FAST RNA-seq Lib Prep Kit (v2) | ABclonal | RK20306 | Reuse | |
| Critical commercial assay | Poly(A) mRNA Capture Module | ABclonal | RK20340 | Reuse | |
| Critical commercial assay | High-range (20 – 1500 mg COD/L) COD kit | Hach | 2565115 | Reuse | |
| Critical commercial assay | QIAamp PowerFecal Pro kit | Qiagen | 51804 | Reuse | |
| Critical commercial assay | Quant-iT™ 1X dsDNA Assay Kit, BR | Life Technologies | Q33267 | Reuse | |

| | | | | | |
|---|---|---|---|---|---|
| Critical commercial assay | Illumina DNA Prep, (M) Tagmentation (96 Samples, IPB) kit | Illumina | 20060059 | Reuse | |
| Dataset | Motor and GI function data | Github/Zenodo | https://doi.org/10.5281/zenodo.17137678 | New | |
| Dataset | Blot images & values | Github/Zenodo | https://doi.org/10.5281/zenodo.17137678 | New | |
| Dataset | Protein assays data | Github/Zenodo | https://doi.org/10.5281/zenodo.17137678 | New | |
| Dataset | Flow cytometry data | Zenodo | https://doi.org/10.5281/zenodo.16929959 | New | |
| Dataset | RNAseq data | SRA | PRJNA1308739 | New | |
| Dataset | Volatile fatty acid analysis data | Github/Zenodo | https://doi.org/10.5281/zenodo.17137678 | New | |
| Dataset | Microbiome sequencing data | SRA | PRJNA1259538 | New | |
| Experimental model: Organism/strain | Thy1-human α-synuclein overexpressing mice "Line 61" | Masliah Lab | N/A | Reuse | Chesselet et al. Neurother 9, 297–314 (2012); Rockenstein et al. J Neurosci Res 68:568–578 (2002) |
| Other | Yeast Casitone Fatty Acids with Carbohydrates (YCFAC) media | Anaerobe Systems | AS-680 | Reuse | |
| Other | Peptone Yeast Extract Broth with Glucose (PYG) media | Anaerobe Systems | AS-822 | Reuse | |
| Other | Mega Medium | N/A | N/A | Reuse | Cheng, A. G. *et al.* Design, construction, and in vivo augmentation of a complex gut microbiome. *Cell* 185, 3617-3636.e19 (2022). |

| | | | | | |
|---|---|---|---|---|---|
| Other | Chopped Meat with Carbohydrates (CMC) | Anaerobe Systems | AS-823 | Reuse | |
| Other | AnaeroPack | MGC | #10-01 | Reuse | |
| Other | Bio-Plex Cell Lysis Kit | Bio-Rad | 171304011 | Reuse | |
| Other | Novex™ Tris-Glycine Mini protein gels ,4-20% | Invitrogen | XP04205BOX | Reuse | |
| Other | Immobilon-P 0.45um PVDF membrane | Merck Millipore Ltd. | IPVH00010 | Reuse | |
| Other | Clarity Western ECL Substrate | Bio-Rad | 170-5061 | Reuse | |
| Other | Nitrocellulose membrane | Thermo Scientific | 88018 | Reuse | |
| Other | Lysing Matrix D Bead beater tubes | MP Biomedicals | 6913500 | Reuse | |
| Other | Hybri-Max - Red blood cell lysing buffer | Sigma Aldrich | R7757 | Reuse | |
| Other | GolgiPlugTM | BD Biosciences | 555029 | Reuse | 1:1000 concentration |
| Other | Fixable Near-IR LIVE/DEAD Stain Kit | Invitrogen | L10119 | Reuse | 1:500 concentration |
| Other | Foxp3/Transcription Factor Fixation/Permeabilization Kit | eBioscience | 00-5521-00 | Reuse | |
| Other | TRIzol reagent | Invitrogen | 15596-026 | Reuse | |
| Other | Sarstedt Z-gel Microtube 1.1ml | Fisher scientific | 50-809-211 | Reuse | |
| Other | Aminex HPX-87H column | Bio-Rad | 1250095 | Reuse | |
| Other | 10 mM VFA mixed standard | Supelco | CRM46975 | Reuse | |

| | | | | |
|---|---|---|---|---|
| Protocol | Beam traversal test | protocols.io | dx.doi.org/10.17504/protocols.io.kxygx4znzl8j/v1 | New |
| Protocol | Pole descent test | protocols.io | dx.doi.org/10.17504/protocols.io.8epv5k9mjv1b/v1 | New |
| Protocol | Wire hang test | protocols.io | dx.doi.org/10.17504/protocols.io.6qpvrw62plmk/v1 | New |
| Protocol | Adhesive removal test | protocols.io | dx.doi.org/10.17504/protocols.io.4r3l21objg1y/v1 | New |
| Protocol | Hindlimb clasping test | protocols.io | dx.doi.org/10.17504/protocols.io.eq2ly4n2mlx9/v1 | Reuse |
| Protocol | Fecal output, score, and water content | protocols.io | dx.doi.org/10.17504/protocols.io.6qpvrw6qplmk/v1 | New |
| Protocol | Total gut transit test | protocols.io | dx.doi.org/10.17504/protocols.io.rm7vz9yq8gx1/v1 | Reuse |
| Protocol | Bead expulsion test | protocols.io | dx.doi.org/10.17504/protocols.io.n92ld6zn7g5b/v1 | New |
| Protocol | Western blot assay | protocols.io | dx.doi.org/10.17504/protocols.io.x54v92z2ql3e/v1 | Reuse |
| Protocol | Dot blot assay | protocols.io | dx.doi.org/10.17504/protocols.io.261gen2xdg47/v1 | Reuse |
| Protocol | Immune cell Isolation | protocols.io | dx.doi.org/10.17504/protocols.io.nm2dc8e | Reuse |
| Protocol | Metagenomics sample processing | protocols.io | dx.doi.org/10.17504/protocols.io.e6nvw15dwlmk/v1 | Reuse |

| | | | | |
|---|---|---|---|---|
| Software/code | FIJI Version 2.10.0 | National Institute of Health (NIH) | https://imagej.net/software/fiji/; RRID: SCR_002285 | Reuse |
| Software/code | FlowJo | BD, Tree Star | https://www.flowjo.com/solutions/flowjo; RRID:SCR_008520 | Reuse |
| Software/code | R code: RNAseq Analysis script | Github/Zenodo | https://github.com/jboktor/fprausnitzii-treatment-rnaseq | New |
| Software/code | GraphPad Prism 10 | GraphPad Software/ Dotmatics | http://www.graphpad.com; RRID:SCR_002798 | Reuse |
| Software/code | R code: Microbiome sequencing analysis script | Github/Zenodo | https://doi.org/10.5281/zenodo.17128213 | New |
| Software/code | STAR aligner v2.7.1 | Github/Zenodo | https://github.com/alexdobin/STAR, RRID:SCR_004463 | Reuse |
| Software/code | DESeq2 package | Github/Zenodo | https://bioconductor.org/packages/release/bioc/html/DESeq2.html, RRID:SCR_015687 | Reuse |
| Software/code | R | R project | v4.3.0; http://www.r-project.org; RRID:SCR_001905 | Reuse |
| Software/code | clusterProfiler package | Bioconductor | http://bioconductor.org/packages/release/bioc/html/clusterProfiler.html; RRID:SCR_016884 | Reuse |
| Software/code | ggplot2 | CRAN | https://cran.r-project.org/web/packages/ggplot2/index.html; RRID:SCR_014601 | Reuse |
| Software/code | MetaPhlAn v4.1.1 | Huttenhower Lab | https://huttenhower.sph.harvard.edu/metaphlan/; RRID:SCR_004915 | Reuse |

| | | | | |
|---|---|---|---|---|
| Software/code | HUMANnN v3.9 | Huttenhower Lab | https://huttenhower.sph.harvard.edu/humann; RRID:SCR_014620 | Reuse |
| Software/code | mia package | Bioconductor | https://bioconductor.org/packages/release/bioc/html/mia.html; RRID:SCR_023619 | Reuse |
| Software/code | “mediation” R package | CRAN | https://cran.r-project.org/web/packages/mediation/index.html; RRID:SCR_026984 | Reuse |
